# Polarized localization of phosphatidylserine in endothelium regulates Kir2.1

**DOI:** 10.1101/2022.08.01.502310

**Authors:** Claire A. Ruddiman, Richard Peckham, Melissa A. Luse, Yen-Lin Chen, Maniselvan Kuppusamy, Bruce Corliss, P. Jordan Hall, Chien-Jung Lin, Shayn M Peirce, Swapnil K. Sonkusare, Robert P. Mecham, Jessica E. Wagenseil, Brant E. Isakson

## Abstract

In the resistance artery endothelium, we show phosphatidylserine (PS) localizes to a specific subpopulation of myoendothelial junctions (MEJs), signaling microdomains that regulate vasodilation. *In silico* data has implied PS may compete with PIP_2_ binding on Kir2.1, a channel involved in vasodilatory signaling. We found 83.33% of Kir2.1-MEJs also contained PS, possibly indicating an interaction where PS regulates Kir2.1. Electrophysiology experiments on HEK cells demonstrate PS blocks PIP_2_ activation of Kir2.1, and addition of exogenous PS blocks PIP_2_-mediated Kir2.1 vasodilation in resistance arteries. Using a mouse model lacking canonical MEJs in resistance arteries (*Eln*^fl/fl^/Cdh5-Cre), PS localization in endothelium was disrupted and PIP_2_ activation of Kir2.1 was significantly increased. Taken together, our data suggests PS enrichment to MEJs inhibits PIP_2_-mediated activation of Kir2.1 to tightly regulate changes in arterial diameter, and demonstrates the intracellular lipid localization within endothelium is an important determinant of vascular function.

## Introduction

Kir2.1 is an inwardly rectifying potassium channel that maintains homeostatic potassium levels inside the cell. The channel is important for regulating cardiovascular homeostasis and is expressed in both the heart and arteries. (*1, 2*) In resistance arteries, active currents have been observed in endothelium but not smooth muscle, (*2*) and global heterozygous knockout of Kir2.1 leads to hypertension in mice. (*1*) In addition to its role in nitric oxide-based flow-mediated vasodilation, (*3*) Kir2.1 is also a contributor to endothelial derived hyperpolarization (EDH), the predominate dilation pathway in resistance arteries. (*2, 4, 5*) Resistance artery endothelium contain a signaling microdomain called the myoendothelial junction (MEJ) which is an endothelial cell (EC) extension through holes in the internal elastic lamina (HIEL) that contacts the underlying smooth muscle cells (SMC). The MEJ is a critical site of heterocellular communication that facilitates EDH vasodilation through gap junctions and localization of proteins, (*6-8*) such as the Ca^2+^-activated intermediate conductance potassium channel (IK_Ca_), and Transient Receptor Potential Cation Channel Subfamily V Member 4 (TRPV4), and alpha hemoglobin. (*4, 9, 10*) While functional data implicates Kir2.1 may be at the MEJ, it has not been demonstrated. Furthermore, Kir2.1 channel function is known to be regulated by lipids, where cholesterol prevents normal activation (*3, 11-14*) and PIP_2_-Kir2.1 interaction lead to channel opening and activation. (*15-19*) Thus, we sought to determine if Kir2.1 was localized to the MEJ and how channel function was regulated by the local lipid environment.

We have previously demonstrated phosphatidylserine (PS) was enriched in MEJs isolated from a vascular cell co-culture (VCCC). (*20*) PS is an anionic phospholipid enriched on the inner leaflet of lipid bilayers and comprises between 3-10% of the lipid composition of a cell. (*21*) Given PS has been implicated in regulating protein localization and inward rectifying potassium channel function (*22, 23*), we sought to determine the contribution of PS to localization and function of Kir2.1 in resistance arteries. While some data indicate PS as a co-activator of Kir2.1 function, (*24, 25*) *in silico* models demonstrate PS can also bind to the PIP_2_ activation site, (*22, 26*) implicating PS may be capable of preventing or competing with the necessary binding of PIP_2_ (*16, 17*) for Kir2.1 channel activity. We therefore hypothesized PS and Kir2.1 localized together at a unique subpopulation of MEJs, where PS could negatively regulate PIP_2_ activation of Kir2.1 and thus modulate the magnitude of vasodilation and total peripheral resistance.

## Results

### Spatial distribution and heterogeneity of MEJs

When arteries are cut longitudinally with the endothelial monolayer facing upwards (*en face* view), potential sites of MEJ formation are detected as holes in the IEL (HIEL; (*4, 27, 28*)). Immunohistochemistry studies on *en face* views have reported heterogeneous protein localization to the HIEL, (*27-35*) with possible juxtaposition to other signaling hubs in the endothelium. To determine if this may be the case, we initially used stitched confocal images of third order mesenteric arteries stained for nuclei, the IEL, and interendothelial junctions (i.e., claudin-5) and visualized approximately 50 fully in view EC per artery (**Fig. 1A**). We found 9.83 HIEL per EC (**Fig. 1A**), and the number of HIEL could be predicted by the size of an EC (**Fig. 1A**), indicating there may be a specific spatial pattern of HIEL. We developed a Matlab program (available online; see **Methods**) to calculate the minimum distance of each individual HIEL to organelles and/or critical signaling hubs within EC and compared it against simulations for a positive (PC), negative (NC), and random (RAND) spatial pattern (**Fig. 1B**). We found a significant difference between real-world and simulated positive and negative values, but no differences between real-world and simulated random values in all tested conditions; nuclei (**Fig. 1C-D**), endoplasmic reticulum (ER) (**Fig. 1E-F**), and interendothelial junctions (**Fig. 1G-H**). The random distribution of HIEL was also evident in first order mesenteric arteries where HIEL density is approximately half of the density in third order arteries (**Fig. S1**), indicating the random localization of the HIEL is not an outcome of high density. Thus, HIELs are randomly distributed with respect to major EC signaling hubs.

**Figure 1:**
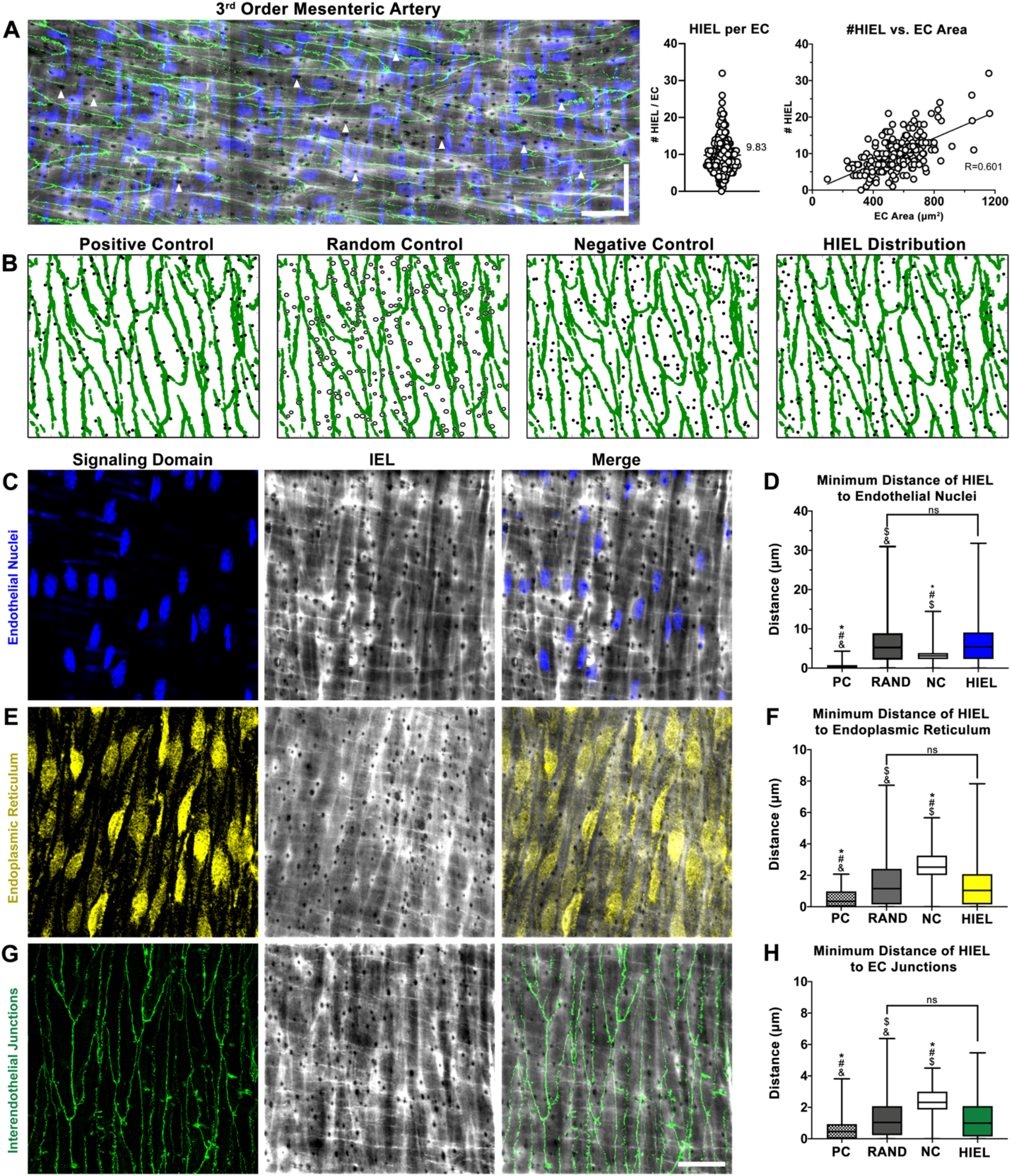
HIEL are randomly distributed with respect to endothelial signaling hubs. **(A)** Representative stitched confocal image of a third order mesenteric artery prepared en face and stained for nuclei (blue) via DAPI, IEL (grey) via Alexa Fluor 488-linked hydrazide, and interendothelial junctions (green) via claudin-5. Scale bar is 30µm in both directions. Quantification of HIEL per EC, and plot of HIEL per EC vs. EC area. N=4 mice, n=4 arteries, n=12 ROIs, and n=205 ECs. (**B**) Graphical outputs of Matlab thresholding and simulations. Interendothelial junction coordinates from *en face* imaged are plotted in green and real-world HIEL centers are plotted in black (HIEL Distribution). Representative outputs of Matlab simulations that generate random (RAND), positive (PC), or negative (NC) HIEL distribution patterns (in black) with respect to interendothelial junctions. Note random simulations also randomly selected an HIEL diameter while PC and NC simulations assumed a uniform HIEL size (circles vs dots, respectively; see **Methods**). (**C**) Representative *en face* confocal image of endothelial nuclei (blue) detected via DAPI and IEL (grey). (**D**) Box and whiskers plot of minimum distance of real-world HIEL centers to endothelial nuclei, compared to Matlab-simulated HIEL centers. N=3 mice, and n=6 arteries, n=18 ROIs, Area=9.75×10^4^µm^2^, and n=1607 HIEL. (**E**) Representative *en face* confocal image of endoplasmic reticulum (yellow) detected via calnexin and IEL (grey). (**F**) Box and whiskers plot of minimum distance of real-world HIEL centers to endoplasmic reticulum compared to Matlab-simulated HIEL centers. N=3 mice, and n=4 arteries, n=9 ROIs, Area=7.96×10^4^µm^2^, and n=1157 HIEL. (**G**) Representative *en face* confocal image of interendothelial junctions (green) detected via claudin-5 and IEL (grey). (**H**) Box and whiskers plot of minimum distance of real-world HIEL centers to interendothelial junctions compared to Matlab-simulated HIEL centers. N=6 mice, and n=10 arteries, n=22 ROIs, Area=1.48×10^5^µm^2^, and n=2166 HIEL. Brown-Forsythe and Welch ANOVA. # is p <0.0001 significant difference to real-world HIEL distribution, * is p <0.0001 significant difference to random simulation distribution, $ is p < 0.0001 significant difference to negative control simulation distribution, and & is p <0.0001 significant difference to positive control simulation distribution.

Given the heterogeneity of proteins observed at the HIEL, (*28-35*) if the HIEL were not specifically localized to EC signaling hubs as demonstrated in **Fig. 1**, another alternative may be not every HIEL contains an EC projection. We used TEM to examine transverse sections of third order mesenteric arteries where cellular projections can be unequivocally identified at the ultrastructural level, thus circumventing the need to rely on an individual protein marker (**Fig. 2A**). We leveraged our *en face* data obtained from our Matlab program and measurements taken from third order mesenteric arteries (**Fig. 2B, Tables S1-2**) to predict how many MEJs would be observed in transverse TEM images (**Fig. S2**; **Tables S1-3**; **Methods**). Based on our calculations, we predicted between 5 and 17.8 HIEL per 1000µm IEL length in the cross-sectional TEM view if all the HIEL were MEJs. Converting the HIEL density in *en face* images to HIEL per 1000µm IEL length falls within this predicted range, validating our prediction (**Fig. 2C**). The measurements taken from TEM cross-sections also fall within this predicted range, where every HIEL contained a cellular projection (**Fig. 2C**); thus, we conclude each HIEL contains a *bona fide* MEJ.

**Figure 2:**
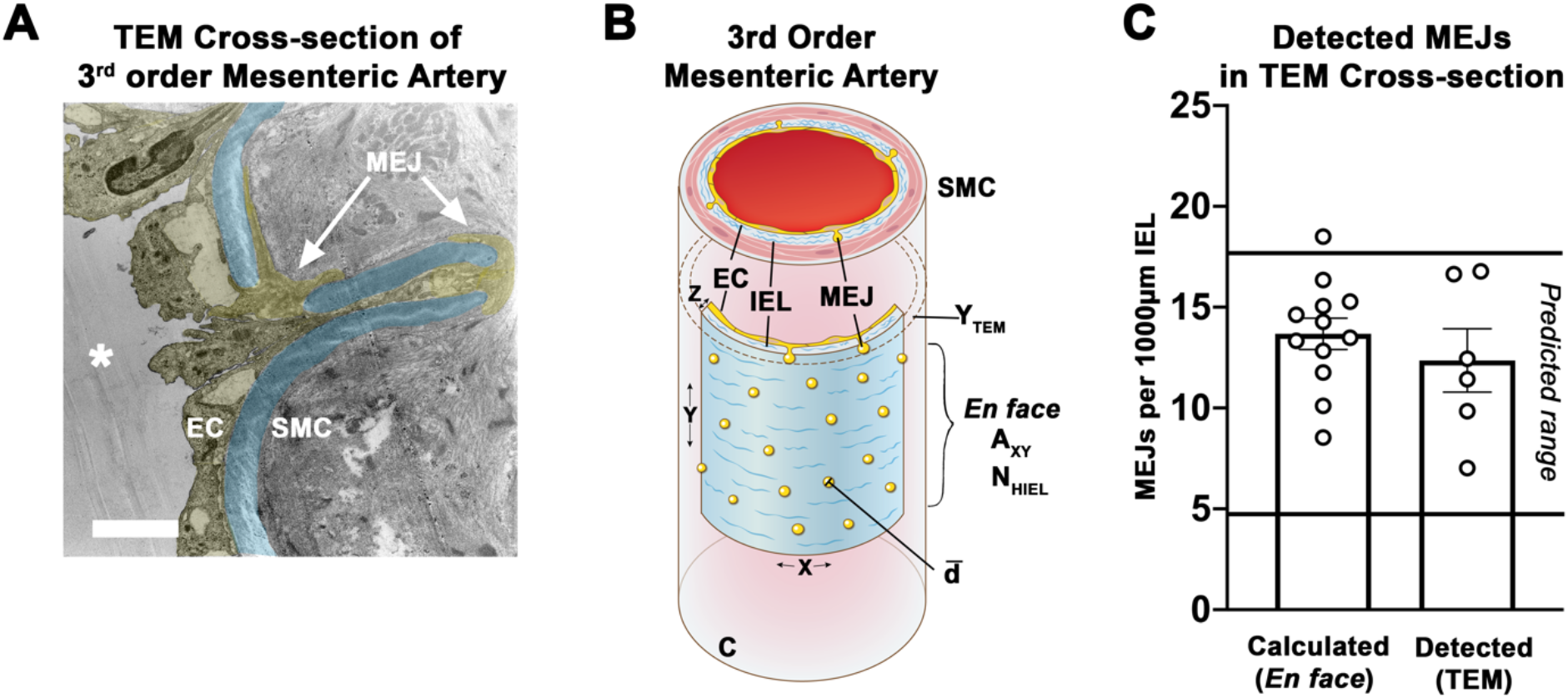
Every HIEL contains an MEJ. (**A**) Transmission electron microscopy (TEM) image of an arterial cross-section at 2K magnification. IEL is pseudo-colored in blue and ECs/MEJs pseudo-colored in yellow. Scale bar 2µm. (**B**) Quantitation of *en face* HIEL from Matlab simulations that were used to predict if every HIEL contains an MEJ (refer to **Methods** and **Tables S1-3**). Where **C** is the circumference of the artery, **d** is the diameter of HIEL, ***Ǡ***_**xy**_ is the area of *en face* images, **N**_**HIEL**_ is the average number of HIEL per image, **Y**_**TEM**_ is the thickness of an individual TEM section, **X** is the width of an *en face* image, **Y** is the height of an *en face* image, and **Z** is the thickness of the arterial wall. (**C**) Quantification of HIEL per 1000µm measured in TEM cross-sections or back calculated from *en face* views, where both values are within the predicted range. For TEM, N=6 mice, n=6 arteries, n=3-5 TEM sections, and 570-964µm IEL length quantified per mouse. For *en face*, N=4 mice, n=4 arteries, n=12 ROIs, and Area=1.67×10^5^µm^2^.

### PS and Kir2.1 define a unique subpopulation of MEJs

If every HIEL contains an MEJ, it was unclear to us what contributes to the heterogenous localization of proteins to the MEJ, especially if there is a random distribution of MEJs with respect to EC signaling hubs. The possibility we considered was the lipid composition of MEJs, due to the emerging evidence of lipid regulation of proteins. (*15, 16, 22, 36*) *In vitro*, we have previously demonstrated enriched PS via lipid mass spectrometry at MEJs isolated from VCCC, ((*6, 20, 37*) **Fig. S3)**), and hypothesized the same lipid accumulation at *in vivo* MEJs. Stitched confocal images of third order mesenteric arteries viewed *en face* revealed PS localization both to the ER where it is synthesized (*38*) and to MEJs in intact arteries (**Fig. 3A-C**). The specificity of the PS antibody was confirmed using a Lactadherin-C2 plasmid in HeLa cells, a protein that specifically binds to PS (**Fig. S3**). (*39-42*) PS was found to be heterogeneously distributed and occupied 13.8% of MEJs (**Fig. 3D**). Analysis of PS-MEJs per EC reveals on average 1.18 PS-MEJs are associated with any individual EC (100% of ECs analyzed, **Fig. 3E**), or 1.95 PS-MEJs if only considering ECs containing at least one (58.04% of all ECs analyzed, **Fig. 3E**). The PS-MEJs did not exhibit a localization pattern within EC (**Fig. 3F**).

**Figure 3:**
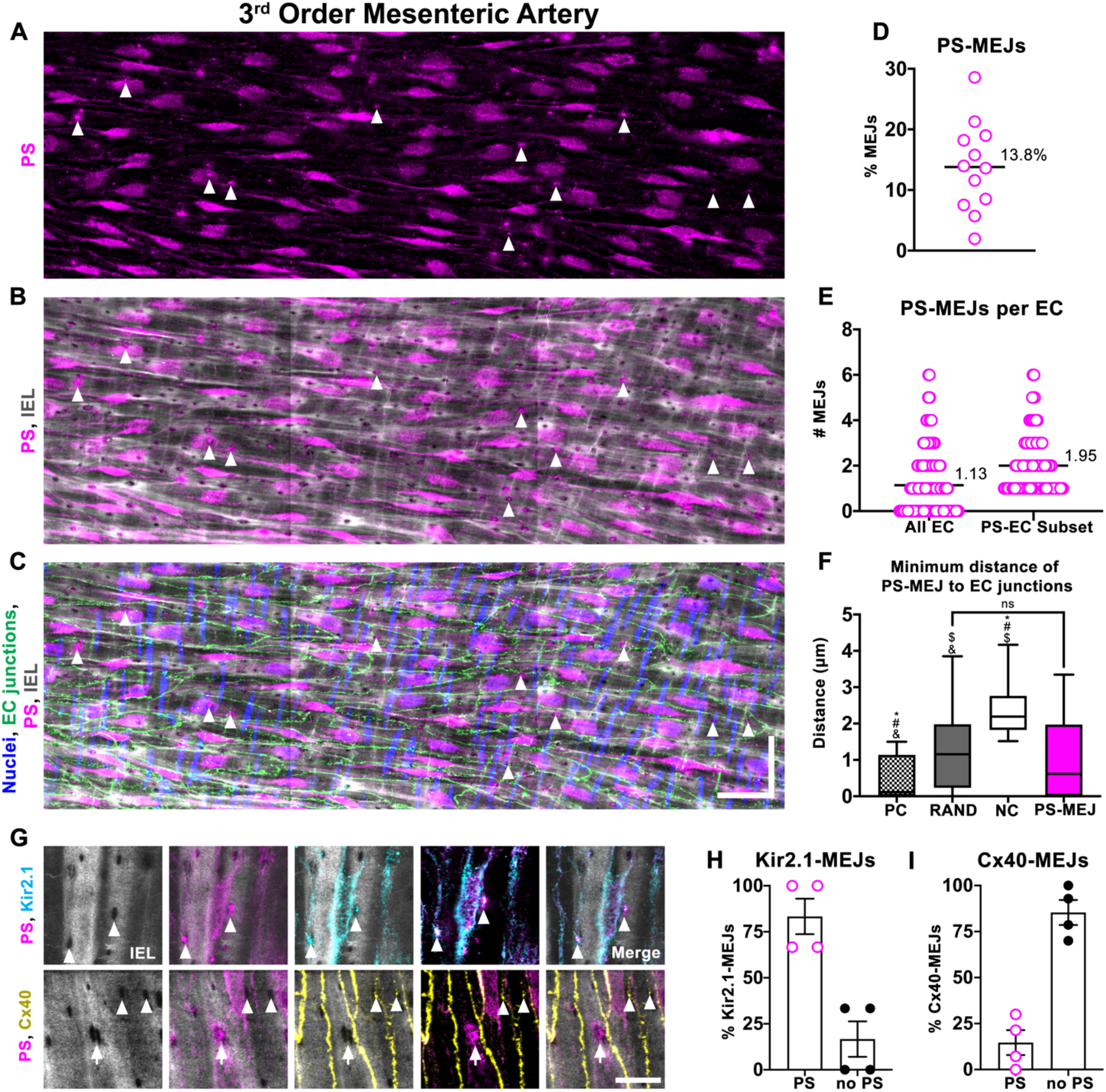
PS colocalizes with Kir2.1 in a subpopulation of MEJs. Representative stitched confocal image of a third order mesenteric artery showing (**A**) PS (magenta), (**B**) PS and IEL (grey) detected via Alexa Fluor 488-linked hydrazide, and (**C**) merge with nuclei (blue) detected via DAPI and interendothelial junctions (green) detected via claudin-5. Arrowheads show PS localized to MEJ. Scale bar is 30µm in both directions. **(D**) Percentage of MEJs in endothelium containing PS. (**E**) Number of MEJs containing PS per EC, where EC borders were defined via claudin-5 staining. N=4 mice, n=4 arteries, n=12 ROIs, and n=205 ECs. (**F**) Spatial analysis of PS-MEJ compared to positive control (PC), random (RAND), and negative control (NC) simulations. Brown-Forsythe and Welch ANOVA with Holm-Sidak multiple comparisons. # is p < 0.0001 significant difference from real-world MEJ distribution, * is p <0.0001 significant difference from random simulation distribution, $ is p <0.0001 significant difference from PC simulation distribution, and & is p <0.0001 significant difference from NC simulation distribution. N=3 mice, n=4 arteries, n=4 ROIs, n=66 PS-MEJs, and Area=4.49×10^4^ µm^2^. (**G**) *En face* images of third order mesenteric arteries with IEL (grey) detected via hydrazide, PS (magenta), Kir2.1 (cyan), and Cx40 (yellow). Scale bar is 10 µm. Arrowheads indicate colocalization of PS and Kir2.1 to MEJ or Cx40 only to MEJ. Arrow indicates localization of PS only to MEJ. (**H**) In-house Matlab analysis to detect incidence of PS and Kir2.1 or (**I**) PS and Cx40 colocalization to MEJ in *en face* images. N=4 mice, n=4 arteries per group.

Because it is well known lipid interactions with ion channels regulate channel activity, (e.g.,(*15, 16, 22, 36*)) we wanted to investigate if PS could have a similar function at the MEJ. I*n silico* data suggests PS has a binding site on Kir2.1, (*22, 26*) an important potassium channel involved in vasodilation, (*2*) demonstrating PS can access and bind the PIP_2_ binding site, possibly interfering with a necessary component of Kir2.1 activation. (*15-19*) Using *en face* immunohistochemistry, we observed 83.33% of Kir2.1-MEJ puncta also contained PS (**Fig. 3G, 3H;** antibody validated in **Fig. S4**). In contrast, localization with PS occurred in only 14.64% of connexin 40 (Cx40)-MEJs, a component of gap junction formation between EC and SMC in arteries (e.g.,(*43*), **Fig. 3I**; antibody validated in **Fig. S5**). Thus, Kir2.1 is highly enriched in PS-MEJs, whereas Cx40 is segregated to non-PS MEJs.

### PS localization to the MEJ dampens PIP_2_ activation of Kir2.1

The strong co-localization of Kir2.1 and PS at MEJs, with *in silico* data indicating a PS binding site, (*22, 26*) led us to hypothesize PS may regulate Kir2.1 function. We wanted to test if PS could regulate Kir2.1 activation in third order mesenteric arteries where Kir2.1 and PS localize to the same MEJs. We used pressure myography to perform dose response curves with NS309, a potent dilator that can activate Kir2.1. (*2*) We observed a reduced dilation to NS309 in the presence of known Kir2.1 inhibitors barium (Ba^2+^) (*44, 45*) at 100µM (**Fig. 4A**) and ML-133 (*46, 47*) at 3.6µM (**Fig. 4B**), verifying previously published results. (*2*) We applied exogenous PS at 10µM to the arteries and observed a similar reduction in NS309-mediated vasodilation (**Fig. 4C**). We confirmed in this experimental setup PS could get to the MEJ (**Fig. S6**). Importantly, the application of PS or known Kir2.1 inhibitors did not influence SMC function (**Fig. 4D-F**; except for Ba^2+^ significantly inhibiting KCl constriction, likely due to off-target effects on SMC Kir6.1). (*48*)

**Figure 4:**
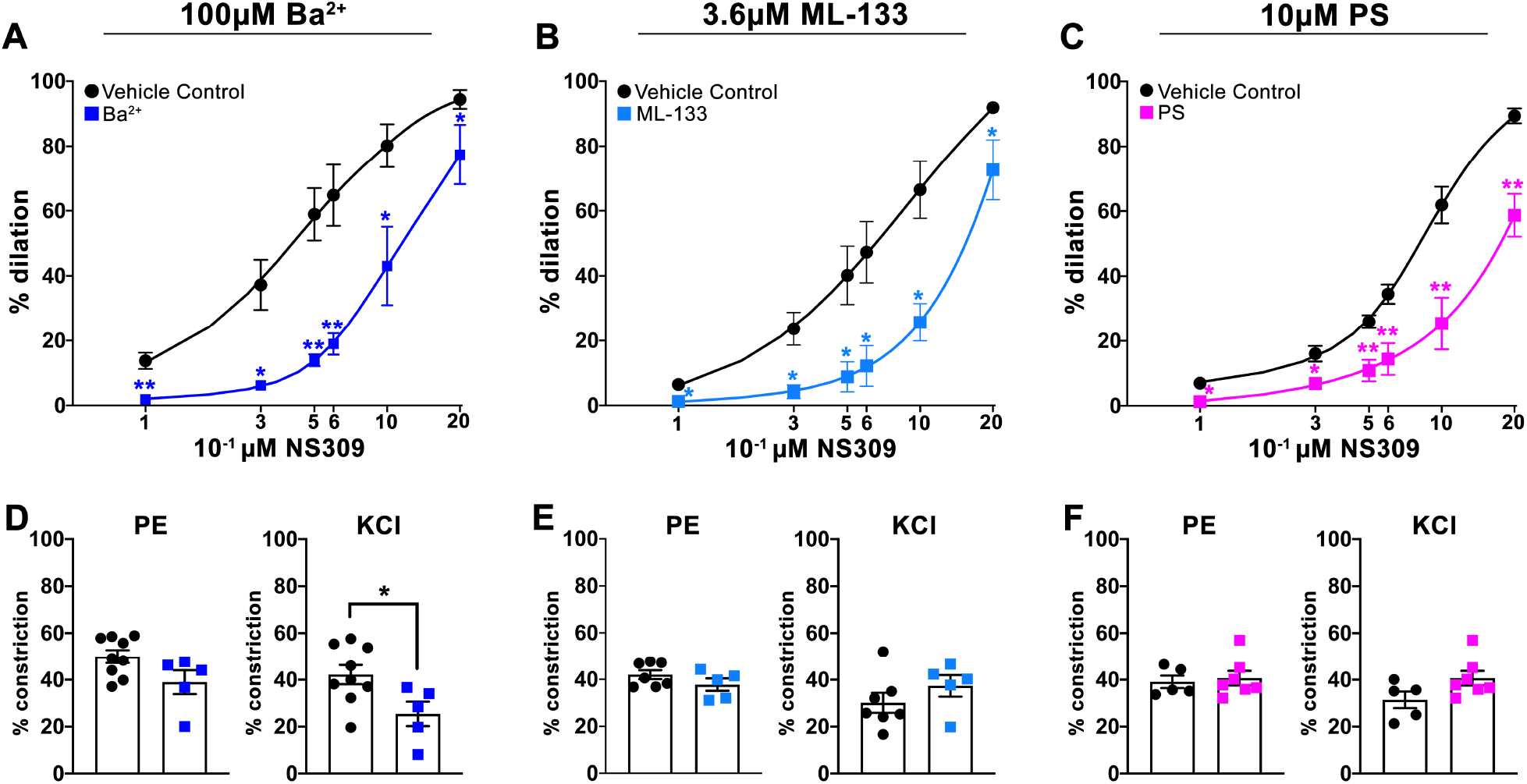
Exogenous PS blocks Kir2.1-mediated vasodilation. Dose response curves to NS309 on third order mesenteric arteries in the presence of (**A**) 100µM Ba^2+^ (**B**) 3.6µM ML-133, or (**C**) 10µM PS compared to vehicle controls of water, DMSO, and ethanol, respectively. (**D-F**) Percent constriction to 1µM PE or 30mM KCl in the presence of each Kir2.1 inhibitor. Student’s t-test, where * p<0.050, and ** p<0.001.

Next, using whole-cell patch clamp electrophysiology and HEK293T cells overexpressing Kir2.1, we tested the direct effect of PIP_2_, PS, or both on whole-cell currents (**Fig. 5A-B**). We observed an increase in Ba^2+^-sensitive currents in both PIP_2_ and PS conditions alone, consistent with previous observations of these lipids contributing to channel opening. (*24, 25*) However, when PS was applied along with PIP_2_, the increase in Ba^2+^-sensitive currents was no longer observed (**Fig. 5A-B**), suggesting the presence of PS can block the well-described PIP_2_-mediated activation of Kir2.1. Because there is strong evidence for PIP_2_-mediated dilation of Kir2.1 in cerebral arteries, (*15*) we sought to test the effect of PS on PIP_2_-mediated dilation on arteries constricted to myogenic tone (**Fig. 5C**). We observed a consistent dilation to 10µM PIP_2_ in pressurized third order mesenteric arteries that was blocked by Kir2.1 inhibitors Ba^2+^ and ML-133 (**Fig. 5D-E**). Surprisingly, PS treatment alone did not influence arterial diameter (**Fig. 5B-C**). In alignment with the electrophysiology data, we found when we pre-incubated the arteries with 10µM exogenous PS for 30 minutes prior to adding PIP_2_, the dilation was significantly decreased (**Fig. 5D-E**); an outcome cogent with PS competing for the PIP_2_ binding site. (*22, 26*) The residual PIP_2_ dilation with a PS preincubation occurs at the same time as PIP_2_ alone, indicating some PIP_2_ is able to access its binding site (**Fig. 5F**), although to a significantly lower extent (**Fig. 5D**). Following a washout period at the end of the experiment, arteries were evaluated for EC function via dilation to 1µM NS309 (**Fig. 5G**) and constriction to 30mM KCl (**Fig. 5H**) to ensure measured differences in PIP_2_ dilation were not due to arteries odf compromised function. The sum of these results indicates Kir2.1 localization to PS-MEJs may be used to tightly regulate vasodilation.

**Figure 5:**
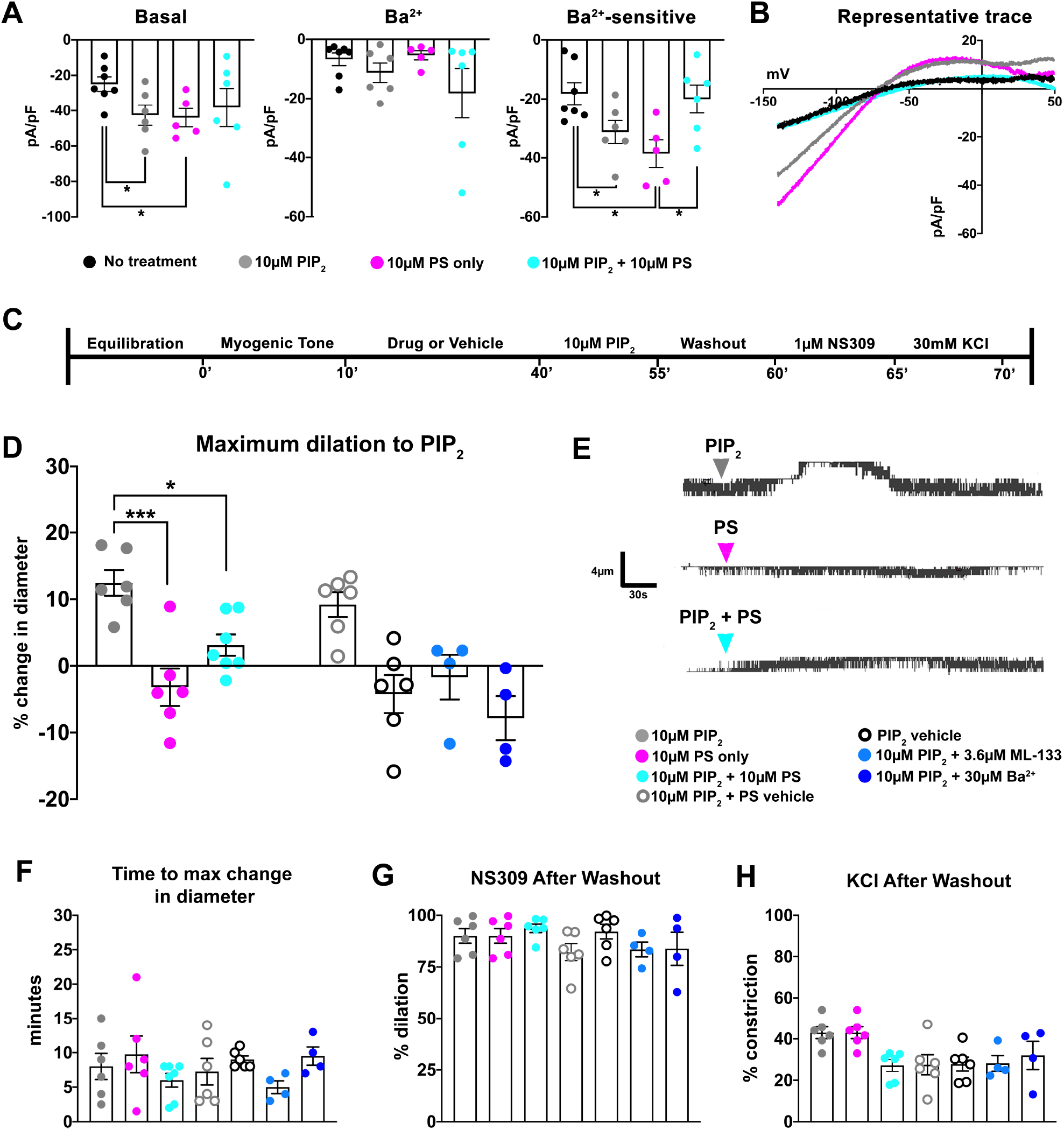
PS inhibits PIP_2_ activation of Kir2.1. (**A**) Average whole cell Kir2.1 currents at - 140mV in transfected HEK293T cells at baseline and with 100µM Ba^2+^. Ba^2+^-sensitive currents were calculated by subtracting Ba^2+^ current from basal current. (**B**) Representative traces of Ba^2+^-sensitive currents. For all graphs: black is basal current, grey is treatment with 10µM PIP_2_, magenta is treatment with 10µM PS, and cyan is treatment with 10µM PIP_2_ + 10µM PS. N=2-3 independent transfections and n=5-7 cells per group. Student’s t-test. * is p<0.050. (**C**) Experimental timeline to assess PIP_2_ dilation in intact arteries. Arteries are equilibrated to temperature and pressure until the development of myogenic tone. After 10 minutes of plateaued diameter, the drug or vehicle treatment was applied to the system for 30 minutes. Next, 10µM PIP_2_ was applied to the system for 15 minutes. The maximum change in diameter within this period was recorded and used for analysis. Following a 5-minute washout, arterial function was assessed via 1µM NS309 and 30mM KCl. (**D**) Maximum change in diameter in each treatment group. N=4-5 mice and n=4-7 arteries per group. Changes in diameter for PS were taken from the same experiments as 10µM PIP_2_ + 10µM PS groups, where the maximum change in diameter was recorded within the 30-minute incubation prior to PIP_2_ treatment. Students t test. * p<0.050 and *** p <0.001. (**E**) Legend for the different groups where (**i**) 10µM PIP_2_ is grey circles, (**ii**) 10µM PS is magenta circles, (**iii**) 10µM PIP_2_ + 10µM PS is cyan circles, (**iv**) 10µM PIP_2_ + PS vehicle is grey open circles, (**v**) PIP_2_ vehicle only is black open circles, (**vi**) 10µM PIP_2_ + 10µM ML-133 is light blue circles, and (**vii**) 10µM PIP_2_ + 30µM Ba^2+^ is dark blue circles. Representative traces of inner diameter from pressure myography experiments demonstrating the effects of 10µM PIP_2_, 10µM PS, or 10µM PIP_2_ + 10µM PS on arterial diameter. (**F**) The timepoint at which maximum change in diameter is achieved for each group. (**G**) The dilation to 1µM NS309 following a 5-minute washout period to assess EC function in each experiment. (**H**) The constriction to 30mM KCl to assess SMC function in each experiment. One-way ANOVA was performed for **F-G**.

### Disrupted PS localization in endothelium increases PIP_2_ activation of Kir2.1

Next, we sought to determine if the MEJ itself was the key driver in PS-Kir2.1 organization, and by extension vasodilation. To do this, we took advantage of floxed elastin mouse (*Eln*^fl/fl^) on an endothelial cell-specific Cre (Cdh5; i.e., *Eln*^fl/fl^/Cre^+^) that had previously been shown to lack IEL in resistance arteries, (*49*) and thus possibly lacked MEJs. The mice had the correct elastin deletion and loss of IEL in resistance arteries (**Fig. 6A-C**; **Fig. S7A-C**), but not large arteries (**Fig. S7D-G**). The disruption of IEL was also evident in the *en face* view with a 56.26% reduction in hydrazide area and the loss of observable, morphologically distinct HIEL (**Fig. 6D-F**). Despite normal tight junction morphology (**Fig. 6E**), the pattern of PS localization in endothelium is drastically changed, where it covers less surface area (**Fig. 6I**), does not accumulate into large puncta (**Fig. 6J**), and has decreased pixel intensity compared to controls (**Fig. 6L-M**).

**Figure 6:**
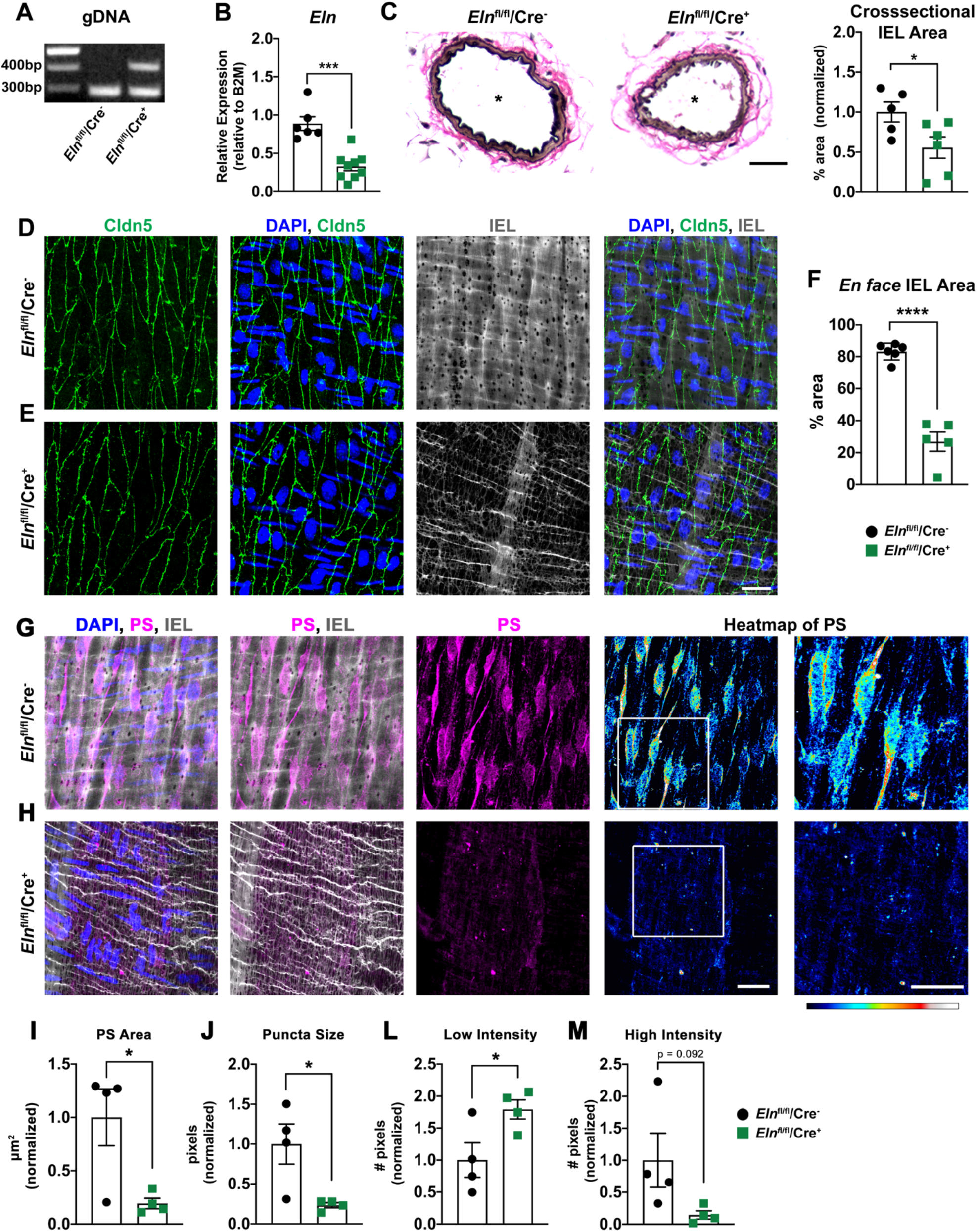
*Eln*^fl/fl^/Cre^+^ mice have disrupted PS localization in endothelium. (**A**) Genomic DNA (gDNA) isolated from EC-rich lung tissue demonstrating the expected excision band of elastin in *Eln*^fl/fl^/Cre^+^ tissue at 410bp. (**B**) qPCR on mesenteric vasculature to assess the mRNA levels of *Eln* in both *Eln*^fl/fl^/Cre^-^ and *Eln*^fl/fl^/Cre^+^ tissue. N=6-9 mice per group. Student’s t-test and *** is p =0.0001. (**C**) Cross-sections of third order mesenteric arteries are stained with Verhoeff elastic stain (black) and counterstained with van Gieson (pink). Percent area with Verhoeff stain is quantified in the graph on the right. * indicates lumen. Scale bar is 30µm. N=5-6 mice per group. Student’s t-test and * is p<0.050. (**D**) *En face* immunohistochemistry on *Eln*^fl/fl^/Cre^-^ and (**E**) *Eln*^fl/fl^/Cre^+^ third order mesenteric arteries where interendothelial junctions (green) are detected via claudin-5, nuclei (blue) are detected via DAPI, and IEL (grey) is detected via and Alexa Fluor 488-linked hydrazide. Scale bar is 30µm. (**F**) Quantification of IEL area in *en face* views, expressed as a percentage of total image area. N=5 mice, n=5-6 arteries. Student’s t-test and **** is p < 0.0001. (**G**) *En face* immunohistochemistry on *Eln*^fl/fl^/Cre^-^ and (**H**) *Eln*^fl/fl^/Cre^+^ third order mesenteric arteries where PS (magenta), nuclei (blue) are detected via DAPI, IEL (grey) is detected via Alexa Fluor 488-linked hydrazide. PS heatmaps were generated using Royal Lookup Tables in ImageJ, where low intensity signal is in cool tones (blue to cyan) and high intensity signal is warm tones (yellow to white). The white box on the heatmap indicates the zoomed in area shown to the right. Scale bars are 10µm. (**I**) Quantification of PS area, (**J**) puncta size, (**L**) low intensity pixels, and (**M**) high intensity pixels in PS *en face* images comparing *Eln*^fl/fl^/Cre^-^ and *Eln*^fl/fl^/Cre^+^ third order mesenteric arteries. N=3-4 mice, and n=4-5 arteries per group. Student’s t-test and * is p<0.050.

In *Eln*^fl/fl^/Cre^+^ mice, Kir2.1 protein and function were unchanged as assessed by quantitative western blot and NS309-mediated dilation, respectively (**Fig. 7A-B**). Given these results, if the PS polarization to MEJs were important for dampening Kir2.1-mediated dilation, application of PIP_2_ would have increased dilation in *Eln*^fl/fl^/Cre^+^ arteries. *Eln*^fl/fl^/Cre^-^ arteries were used at 60mmHg based on their myogenic tone (**Fig. 7C, Fig. S8**). This dilation was not different compared to arteries from C57BL6 mice at 60mmHg (**Fig. 7C**). *Eln*^fl/fl^/Cre^+^ arteries exhibited a significant increase in PIP_2_-mediated vasodilation which was inhibited with addition of 10µM PS or 3.6µM ML-133 (**Fig. 7C**). Interestingly, arteries from *Eln*^fl/fl^/Cre^+^ had an increased time to maximum change in diameter (**Fig. 7D-E**) compared to *Eln*^fl/fl^/Cre^-^ or C67Bl6 arteries, which is reflective of a continued dilation in *Eln*^fl/fl^/Cre^+^ arteries compared to a transient dilation in control experiments. A continued dilation coincides with PS not being spatially oriented to negatively regulate the PIP_2_-mediated activation of Kir2.1 in *Eln*^fl/fl^/Cre^+^ arteries, thus resulting in an uncontrolled dilation. After a washout period, EC and SMC function were evaluated with 1µM NS309 and 30mM KCl respectively (**Fig. 7F-G**).

**Figure 7:**
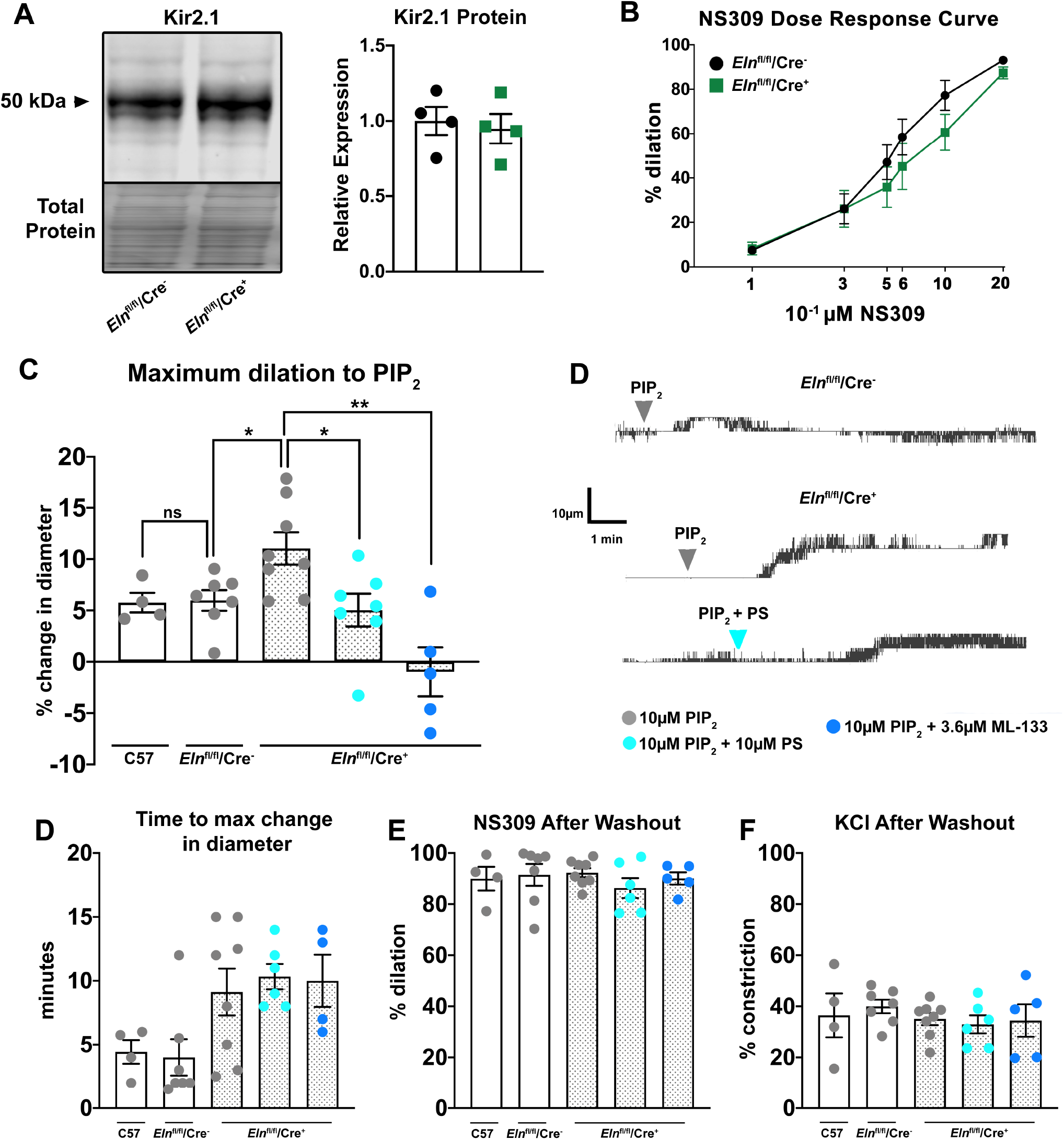
*Eln*^fl/fl^/Cre^+^ mice have increased dilation to PIP_2_. (**A**) Western blot on mesenteric vasculature to detect Kir2.1. Total protein was used as a loading control. Quantification was performed as a percent of total protein, then normalized with respect to the average protein expression in *Eln*^fl/fl^/Cre^-^ samples. N=4 mice per group. Student’s t-test. (**B**) NS309 dose response curve on *Eln*^fl/fl^/Cre^-^ and *Eln*^fl/fl^/Cre^+^ third order mesenteric arteries. N=4 mice and n=8-11 arteries per group. Student’s t-test was performed for each dose. (**C**) Dilation to 10µM PIP_2_ at 60mmHg. N=4-5 mice, n=4-8 arteries per group. Student’s t-test. * is p<0.050 and ** is p <0.010. (**D**) Representative traces from pressure myography experiments demonstrating the dilation to PIP_2_ in *Eln*^fl/fl^/Cre^-^ arteries and *Eln*^fl/fl^/Cre^+^ arteries with or without a 30-minute PS preincubation. (**E**) The timepoint at which maximum change in diameter was achieved for each group. (**F**) Dilation to 1µM NS309 following a 5-minute washout period to assess EC function in each experiment. (**G**) Constriction to 30mM KCl to assess SMC function in each experiment.

## Discussion

### Subpopulations of MEJs have distinct functions

Here we have demonstrated PS defines a functionally distinct subpopulation of MEJs where it localizes with and negatively regulates Kir2.1 through preventing PIP_2_ activation and is spatially separated from MEJs containing the gap junction protein Cx40 (**Fig. 8**). It is likely the primary function of PS-Kir2.1 MEJs is to generate vasodilatory hyperpolarization, which can be communicated to SMCs via nearby gap junction-MEJs. Segregating the generators and communicators of hyperpolarization to distinct subpopulations of MEJs (**Fig. 8**) represents an important mechanism to tightly regulate resistance arterial diameter. This observation, coupled with the functional data herein, strongly suggests heterogenous MEJ populations may exist to carry out different functions, either direct heterocellular communication or to function in a vasodilatory capacity. Our data defines a unique MEJ subpopulation, suggesting a role for PS in modulating PIP_2_ activation of Kir2.1 within the artery.

**Figure 8:**
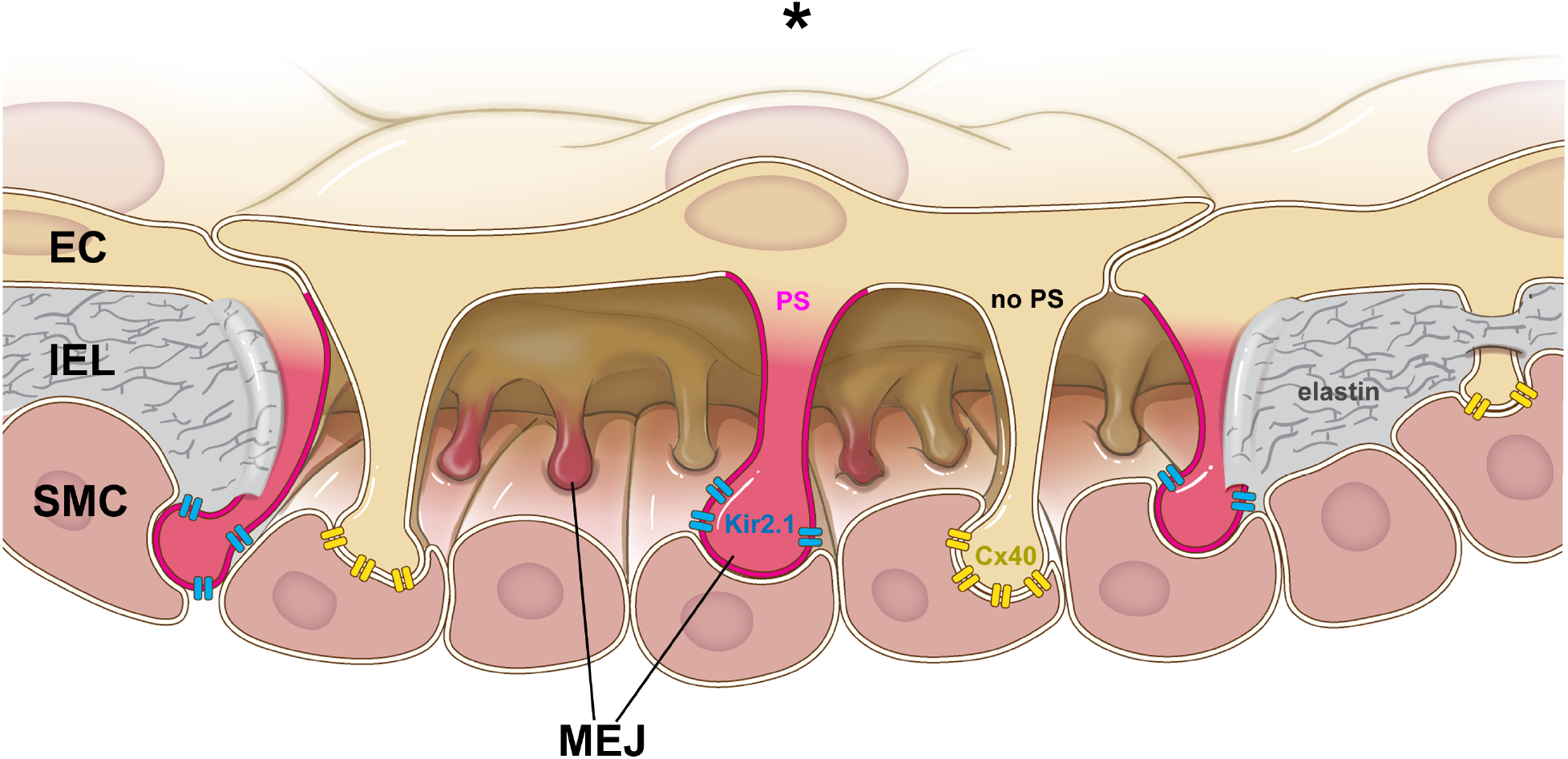
Heterogeneous lipid localization to MEJs results in functionally distinct subpopulations. A zoomed in look at MEJs in the arterial wall. ECs are at the top, SMCs are at the bottom, MEJs are between the two and are encased in IEL proteins such as elastin (grey). Kir2.1 channels (blue) localize to MEJs enriched with PS (magenta), while the gap junction protein Cx40 (yellow) localizes to MEJs that do not contain PS (beige). Image demonstrates subpopulations of MEJs exist within the artery and implies they have unique, specific functions. For clarity, the IEL is only shown at the edges of the image. * is lumen.

Another MEJ population that has been described is the a-kinase anchoring protein 150 (AKAP150)-TRPV4 MEJ, where AKAP150 is required for adequate activation of TRPV4 activation and vasodilation. (*9*) This co-localization at the MEJ is important for facilitating the EDH pathway. (*10*) In a separate study on rat resistance arteries, Cx37 and the IK_Ca_ colocalize in a subpopulation of MEJ, suggesting some potential for potassium channel and gap junction MEJ overlap. (*30*) Further spatial and functional analysis of the Cx37-IK_Ca_ would be necessary to fully understand the role for the co-localization, but we speculate the Cx37-IK_Ca_ MEJ population identified in rat resistance arteries may exist in small groups of ECs and correlate with heterogenous responses of ECs to receptor agonism. (*50*) Interestingly, the Cx37-IK_Ca_ population contrasts with the spatial separation of Kir2.1 and Cx40 we demonstrate here. However, it is not surprising there is a difference in colocalization with gap junctional MEJs between these two potassium channels because of the differences in regulation, activation, and potassium flux. It is unknown what percentage of total MEJs the AKAP150-TRPV4 or Cx37-IK_Ca_ populations occupy. The data presented in this paper includes rigorous quantification of PS localization to MEJs and evaluation of spatial pattern within the endothelium. Future efforts to understand MEJ heterogeneity should use similar quantification methods for understanding possible coordinated function of unique subpopulations.

In our *en face* immunohistochemistry PS localizes to the perinuclear region (**Fig. 3**) suggesting localization in the ER. Given the distinct subpopulation of MEJs that PS occupies, there is likely a specific mechanism of PS transport to facilitate accumulation in the MEJ. Energy-driven lipid transport within a cell occurs via specific lipid transport proteins (LTP) at membrane contact sites, where two lipids are transported in opposite directions. A family of LTPs, called oxysterol-binding proteins (ORPs), have been implicated in the trafficking of PS to the plasma membrane. (*51-53*) It could be the expression of an ORP in ECs drives deposition of PS to specific the MEJs. Kir2.1 trafficking through the ER is also well described, and indeed, we see perinuclear staining in Kir2.1 *en face* preparations (**Fig. 3G**). (*54, 55*) Given the specific, targeted localization of Kir2.1 to PS-enriched MEJs, we speculate PS may also function in trafficking Kir2.1 to these signaling microdomains.

### Phosphatidylserine regulation of Kir2.1

Kir2.1 channel function is dependent on the presence of both PIP_2_ and an anionic phospholipid. (*24, 25*) We have reproduced this data and shown both PS and PIP_2_ can activate the channel in an electrophysiology experiment. However, when added together, the lipids did not have an additive effect on channel activity, but rather did not increase current beyond baseline (**Fig. 5A-B**). In agreement with this finding, our pressure myography data shows addition of exogenous PS blocks PIP_2_ activation of Kir2.1, suggesting the role of PS at the MEJ may be primarily to inhibit Kir2.1 channel activity as opposed to activating the channel. Our data also demonstrates exogenous application of PS inhibits agonist-mediated Kir2.1 activation in intact arteries, also reinforcing the idea of PS as a negative regulator in this tissue. Since flux through the Kir2.1 channel hyperpolarizes the SMC membrane and induces relaxation, PS localization with Kir2.1 at the MEJ may dampen the K^+^ flux through Kir2.1 to prevent overshooting a dilatory effect.

It has been demonstrated relative concentrations of lipids can have differential effects on channel function (*24*), and in MEJs where PS is localized, it is likely the PS concentration is magnitudes higher than the local PIP_2_ concentration, such that PIP_2_ interactions with the channel are limited. Although the local concentration of PIP_2_ remains elusive, we speculate that PIP_2_ has a short half-life at the MEJ given the importance of the PIP_2_ cleavage product, inositol-1,4,5-trisphosphate, in facilitating heterocellular signaling to the SMC and the enrichment of the second cleavage product, diacylglycerol, via lipid mass spectrometry. (*20, 56*) Thus, local PS enrichment at the MEJ may be antagonizing the PIP_2_ requirement for channel opening, and may explain why in intact arteries PS blocks PIP_2_ activation of Kir2.1. Together with the pressure myography results, it is evident the balance between PS and PIP_2_ within the membrane can have differential effects on the protein. We conclude a relatively higher PS concentration at the MEJ functions to negatively regulate Kir2.1 opening.

Our pressure myography data indicates exogenous PS alone does not influence arterial diameter, which is unexpected when compared to the electrophysiology experiments where PS activates the channel. A possible reason for the differential results of PS influence on Kir2.1 could be the differences in cellular lipid composition of HEK293T cells and intact arteries. (*24*) For example, the plasma membrane of ECs in intact arteries is complex, with signaling microdomains defined by protein and lipid composition. One example of this is the high density of caveolae in endothelium. (*57*) Caveolae are crucial for plasma membrane organization and are also enriched with cholesterol, with some studies also implicating specific localization of PS and PIP_2_ to caveolae. (*58, 59*) It is possible there is interplay between PS and cholesterol, a known depressor of Kir2.1 activity (*3, 11, 12*), in intact arteries that prevents PS alone from having a dilatory effect. Indeed, this could be another mechanism by which cholesterol prevents Kir2.1 activation. While we don’t specifically test if the PS-Kir2.1 MEJ population also contains caveolae, it is highly likely caveolae are present within this population given the high density of caveolae in the endothelium and at MEJs. (*57, 60, 61*) In the future, the interplay between cholesterol, PS, and PIP_2_ at the MEJ should be investigated in the context of Kir2.1 and other ion channels.

### The role of MEJ in facilitating phosphatidylserine regulation of Kir2.1

We demonstrate the inhibitory effect of PS on PIP_2_ activation using arteries from a mouse model lacking typical HIEL morphology and thus MEJ formation (*Eln*^fl/fl^/Cre^+^). In this mouse, the PS localization within the intact resistance arterial endothelium is completely disrupted (**Fig. 6**), covering less surface area and demonstrating reduced accumulation into punctate (reduced high intensity signal). Notably, Kir2.1 protein expression is unchanged, yet *Eln*^fl/fl^/Cre^+^ arteries exhibited an increased dilation to PIP_2_, which was brought back down to control levels by adding PS to the artery. These results strongly support our findings of PS negatively regulating PIP_2_-mediated Kir2.1 vasodilation. Based on the result that the PIP_2_ dilation can be recovered to control levels with the addition of PS or ML-133 (a known Kir2.1 inhibitor), (*47*) we attribute the increase dilation to the increased access of PIP_2_ to Kir2.1. This increased dilation is likely due to the disrupted PS localization in the endothelium rather than a direct result of reduced elastin protein, which is cogent with Kir2.1 not demonstrating any known adhesion properties or being regulated by ECM proteins. However, there are some studies linking integrin signaling to ion channel function (*62*) and in particular, increasing current through Kir2.1. (*63*) In *Eln*^fl/fl^/Cre^+^ resistance arteries, integrin signaling is likely decreased since there is a reduction of integrin substrate (ECM proteins). Thus, integrins are an unlikely culprit because of an observed increase in PIP_2_-mediated dilation.

### Limitations of our study

Our interpretation of PS negatively regulating PIP_2_-mediated activation assumes local PIP_2_ concentration at the MEJ is orders of magnitude lower compared to PS. It is a technical challenge to establish the local lipid concentration of PIP_2_ at the MEJ; it was not detected in lipid mass spectrometry of *in vitro* MEJs, but rather, a product of its cleavage, DAG, was found to be enriched at MEJ compared to EC or SMC. (*20*) This suggests to us the half-life of PIP_2_ at the MEJ is short lived because its cleavage products are needed for downstream vasodilatory signaling. (*20, 56*) Even though we are unable to quantify the exact concentration of PIP_2_ at MEJs, our lipid mass spectrometry (*20*) and *en face* evidence (**Fig. 3**) demonstrate a polarized enrichment of PS to a subpopulation of MEJs, suggesting an orders of magnitude higher concentration of PS at these MEJs compared to other lipids typically in the plasma membrane, including PIP_2_; thus reducing relative concentrations of these other lipids at this signaling domain. This paired with a conceivably short PIP_2_ half-life at the MEJ strongly suggests a minute local concentration of PIP_2_ and implies Kir2.1 is surrounded by predominately PS at these signaling microdomains.

We present evidence of lipid regulation of Kir2.1 using an *Eln*^fl/fl^/Cre^+^ mouse model. It should be noted this is the first study to investigate endothelial function on these mice. While ECs appear to function normally (**Fig. 7B**), their myogenic tone (**Fig. S8**) is significantly altered at higher pressures with deviation from controls beginning at 80mmHg, a pressure within normal physiological range. However, it is clear that the polarized localization of PS is lost in these arteries, and despite Kir2.1 protein expression and basal function remaining unchanged (**Fig. 7A-B**), the sensitivity to PIP_2_-mediated vasodilation is increased (**Fig. 7C**), strongly supporting our evidence of MEJ-localized PS regulating Kir2.1 function. Thus, the increased sensitivity to PIP_2_ dilation is unlikely the result of altered myogenic tone.

### Outlook

Our data demonstrates how spatial polarization of a lipid in native ECs can regulate the function of an ion channel, where channel activation by one lipid is modulated by the concentration of a second lipid. While lipid regulation of ion channels and proteins is well accepted, homogenous lipid distributions across the plasma membrane are often assumed, such as in simulations or liposome experiments, and are not representative of native lipid environments. Here we have demonstrated for the first time how intracellular spatial polarization of a lipid within intact tissue can selectively regulate the function of an ion channel for essential physiological function. Understanding the plasma membrane composition surrounding ion channels or proteins in their physiological environment could clarify results from lipid-protein interaction simulations, identify new lipid regulators, or explain tissue-specific function. Thus, investigation of lipid microenvironments in cells may offer a mechanism for differential ion channel function across tissues. Although more work needs to be done to understand how lipid dynamics affect ion channel function, the data presented herein provide a foundation for understanding ion channel regulation by compartmentalization of plasma membrane lipids.

## Materials and Methods

### Mice

The elastin gene, *Eln*, was selectively knocked out of ECs via a VE-cadherin cre (*Eln*^fl/fl^/Cdh5Cre^+^). These mice were on a C57Bl/6 background, and both sexes within the age range of 10-20 weeks were used for experiments. For experiments on this knockout mouse, *Eln*^fl/fl^/Cdh5Cre^-^ littermates were used. For all other experiments, C57Bl6/J male mice between 10-20 weeks were used. Mice were fed a normal chow and housed under a 12-hour light/dark cycle. Mice were sacrificed via CO_2_ inhalation with a secondary method of cervical dislocation. All experiments were approved by the University of Virginia Animal Care and Use Committee.

### Antibodies and plasmids

The following connexin plasmids were a kind gift of Janis Burt: pcDNA3.1 hygro mouse Cx37, pcDNA3.1 puro mouse Cx40, and pcDNA3 neo mouse Cx43. We purchased pEGFP-C1 neo Lactadherin-C2 (Addgene, 22852) and pCAG-Kir2.1-T2A-tdTomato (Addgene, 60598). pcDNA3.1 Panx1 was cloned in house.

Antibodies used were rabbit anti phosphatidylserine (Biomatik, CA30389), mouse anti KCNJ2 (Sigma, SAB5200027), mouse anti connexin 40 (Thermofisher, 37-890), goat anti calnexin (Abcam, ab219644), and Alexa-Fluor linked hydrazide 488 (Thermofisher, A10436) or 647 (Thermofisher, A20502).

### *En face* immunohistochemistry

Mesenteric arteries were fixed with 4% paraformaldehyde at 4°C for 15 minutes. Following PBS washes, arteries were cut *en face* using microscissors and pinned out with tungsten wire (Electron Tube Store, 1439, 0.013mm diameter) on small Sylgard squares (Electron Microscopy Sciences, 24236-10, dimensions: ∼0.5mm thick x ∼1cm tall x ∼0.5cm wide). Next, *en face* preps were permeabilized in 0.2% NP40/PBS at RT for 30 minutes, blocked with BSA or animal serum for 1 hour, and incubated with primary antibody 1:100 at 4°C overnight in blocking solution. Blocking solutions were either 5% serum or 10% BSA (for PS experiments) in 0.2% NP40/PBS.

The next day *en face* preps were washed in PBS and incubated with secondary antibodies (1:400) and hydrazide at RT for 1 hour in blocking solution. Alexa Fluor 488- or 647-linked hydrazide (1mM in diH_2_O stored at 4°C for up to 6 months) and diluted 1:1250-1:2500 in secondary antibody solution to visualize IEL. Next, the arteries were washed 3x 10 minutes in PBS. The third PBS wash included DAPI (Invitrogen, D1306) at 1:5000 from a 5mg/ml stock solution. Following PBS washes, the Sylgard square was lifted out of the plate, the back dried with a Kimwipe, and then placed onto a microscope slide. A droplet of Prolong Gold with DAPI was dispensed on top of the Sylgard square (∼20µl, Invitrogen, P36931). Lab tape was used to secure a square coverslip to the microscope slide prior to imaging. Images were obtained on either a Zeiss 880 LSM with Airyscan (Matlab analysis images) or Olympus Fluoview 1000 with a 40x oil objective and 1.8 zoom.

### Analysis of stitched confocal *en face* images

Stitched confocal images of arteries prepared *en face* were taken as three images on a Zeiss 880 LSM confocal microscope with a 40x objective and 1.8 zoom. Each individual image was analyzed using our in-house Matlab analysis program in order to identify HIEL and PS-HIEL. Next, individual ECs were traced using claudin-5 staining in Image J and assigned a number. Individual HIEL data obtained from Matlab was then manually organized by EC number in Microsoft Excel for analysis.

### Custom analysis of immunofluorescence *en face* images

Briefly, each immunofluorescence channel of an *en face* image was subject to customized thresholding in order to detect HIEL and EC signaling hub (nuclei, interendothelial junctions, or endoplasmic reticulum depending on experiment). The brightness and contrast settings were adjusted around difficult-to-detect HIEL in ImageJ prior to thresholding in Matlab. For claudin-5 experiments (not calnexin or nuclei), the ImageJ line tool was used to assist in detection of low-intensity claudin-5 signal at interendothelial junctions, and nonspecific background noise in the middle of the cell was removed to facilitate correct thresholding in Matlab program. Thresholding was confirmed as accurate through a visual output with thresholded area overlayed on original Z-stack image.

This thresholding was used to calculate the minimum distance of HIEL center points to the EC signaling hub considered and this distribution was plotted as a box and whiskers plot. The distribution was then compared to positive (PC), negative (NC), and random control (RAND, via Monte Carlo) simulations in order to determine the spatial distribution. PC simulations generate HIEL within 1µm of signaling hub, NC generate HIEL between 1.5-4.5µm of signaling hub, and RAND simulations generate randomly distributed HIEL. For PC and NC simulations, each HIEL simulated was the average diameter of all HIEL within the image. Representative plots in **Fig. 1B** show HIEL center points of equal diameter. For RAND simulations, in addition to a random position being selected for the HIEL, a random HIEL diameter was chosen within the range of HIEL sizes measured for each image. Representative plot in **Fig. 1B** shows RAND simulated HIEL of varying diameter.

Puncta detection of PS, Kir2.1, and Cx40 was also done in this automated Matlab program, where a puncta was defined as being within an HIEL if its center point was within 0.75µm of the HIEL center point. This definition was verified as accurate via a visual output from the program. The same PC, NC, and RAND simulations were run on PS-MEJs to determine spatial patterns. Data was transferred to GraphPrism, plotted as a box and whisker plot, and Brown-Forsythe and Welch ANOVA and Holm Sidak’s multiple comparisons statistical tests were performed to detect differences. Outliers were removed (ROUT Q=1%). N values for HIEL in figure legends include entire set and do not reflect data points removed via outlier analysis. In-house Matlab code, example images, and instructions on how to use the code are available on Github: https://github.com/claireruddiman/Spatial-Distribution.

### Transmission Electron Microscopy

Third order mesenteric arteries were dissected from mice and fixed in 4% paraformaldehyde / 2.5% glutaraldehyde / PBS for a minimum of 4 hours. Arteries were then processed at the UVA Advanced Microscopy Facility. The arteries were washed in cacodylate buffer, incubated in 2% osmium tetroxide for 1 hour, washed in cacodylate buffer, then dehydrated with ethanol washes prior to incubation in 1:1 propylene oxide:epoxy resin overnight. The next day the samples were incubated in a 1:2 PO/EPON mixture for 2 hours, then a 1:4 PO/EPON mixture for 4 hours, then 100% EPON overnight. The next day, the samples were baked in an oven at 65°C prior to sectioning. Ultrathin 70nm sections were mounted on a mesh copper grid. Sections were contrast stained with 0.25% lead citrate for 5 minutes, 2% uranyl acetate for 20 minutes, and then again 0.25% lead citrate for 5 minutes. Sections were visualized and imaged using a JEOL 1230 Transmission Electron Microscope. To avoid the possibility of double counting an HIEL, only 1 section per 5µm artery length was analyzed, and a minimum of 500µm of IEL length was analyzed per artery.

### Predicting the incidence of myoendothelial junctions in transmission electron microscopy cross-sections

First, we considered basic descriptive data obtained from *en face* images (**Supp. Table 1**) including the circumference of the artery (**C, Supp. Fig. 2A**) the average diameter of HIEL (**d, Supp. Fig. 2B**), and spatial density of HIEL (**ρ**_**HIEL**_, **Supp. Fig. 2C**), which was calculated using **Eq. 1**.

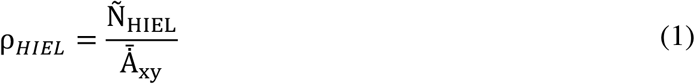

**Ñ**_**HIEL**_ and **d** were quantified via the in-house Matlab program, and **C** was measured as the width of the artery when prepared *en face*. For this calculation, we assume the measured circumference of the artery is equivalent to the circumference of the IEL (**C**_**IEL**_).

Next, we considered the transverse view of the artery when prepared for TEM imaging (**Supp. Table 2**). The TEM section is a thin cross section sliced from a third order mesenteric artery where the thickness of an individual section is **Y**_**TEM**_. Since our quantification of HIEL thus far is from the *en face* view, we wanted to quantify the IEL *en face* surface area in 1 TEM section in order to predict the density of HIEL in TEM sections. A single TEM cross-section corresponds to an *en face* surface area of 20.902µm^2^, which was calculated using **Eq. 2**.

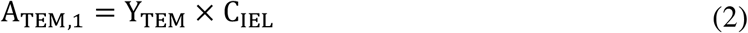

An important note is **d** is much larger than **Y**_**TEM**_, such that the average HIEL will span 30 TEM sections **Eq. 3**.

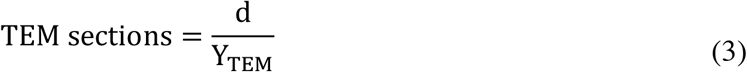

For conceptual clarity, we will consider 6 TEM sections equally spaced apart along artery length **d**, which corresponds to an IEL surface area of 627.06µm^2^ **Eq. 4** and an IEL length of 1791.6µm **Eq. 5**.

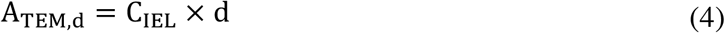

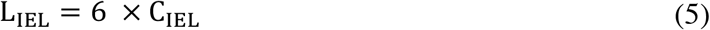

Using **ρ**_**HIEL**_ and **A**_**TEM, d**_, we calculated we would detect a total of <u>9 unique HIEL across a sectioned artery length of</u> **<u>d</u>** with **Eq. 6**.

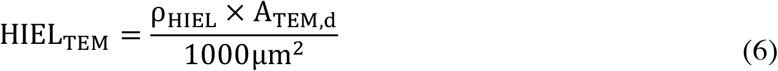

The final variable we considered in our prediction is the spatial distribution of holes throughout the IEL. For this, we considered three cases of distribution (**Supp. Table 3**): (1) uniform distribution (maximum detection), (2) random distribution, and (3) sparse distribution (minimum detection).

For Case 1, we considered 6 TEM sections over artery length **d** and assumed each of the 9 HIEL were exactly aligned with the start and end of the sectioned area (**HIEL**_**TEM**,**6E**_) such that the <u>HIEL distribution is uniform</u>, and the number of HIEL detections by the microscope user is 54 (**Eq. 7**).

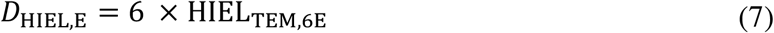

However, due to the nature of TEM, the entire cross-sectional area cannot be imaged due to visibility blockages by grid lines, indicating all 54 HIEL cannot be imaged by the user, even if they were present. In order to account for this technical challenge, the predicted number of HIEL detections can be normalized to the length of IEL in the TEM image. This normalization process results in a numerical value with the units of HIEL per length of IEL. **Thus, if HIEL are uniformly distributed, then we expect 30 HIEL per 1000µm IEL length** (**Eq. 8**).

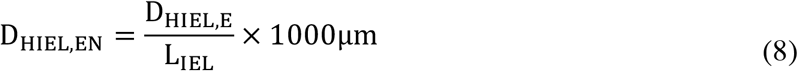

However, based on the Matlab simulations (**Figure 1**), we have shown HIEL are randomly distributed. We therefore incorporated a random distribution of HIEL into our prediction. Thus, instead of assuming each HIEL would have 6 detections across each of the 6 sections, we assumed each HIEL had a different number of detections due to a random alignment with the beginning and end of the length sectioned area. The random distribution of detections was considered as follows: 2 HIEL with 6 detections each (12 detections), 1 HIEL with 5 detections each (5 detections), 2 HIEL with 4 detections each (8 detections), 1 HIEL with 3 detections each (3 detections), 1 HIEL with 2 detections each (2 detections), and 2 HIEL with 1 detection each (2 detections), bringing the total to 32 detections (D_HIEL,R,_ adjusted **Eq. 7** random distribution). Using the same normalization process as described above, we determined **if HIEL are randomly distributed, we expect 17.8 HIEL per 1000µm length of IEL** (**Eq. 9**).

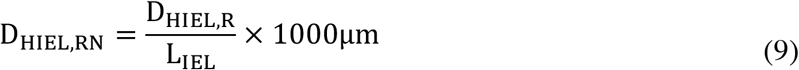

Due to the random distribution of HIEL, there are some areas of IEL with relatively few HIEL. Thus, a third case is considered to reflect a sparse distribution scenario. In this minimum case scenario, each of the 9 HIEL are only aligned with 1 TEM section and thus is only detected once during imaging. For 9 HIEL considered across 6 sections, with each HIEL appearing in only 1 section, there are 9 total detections (D_HIEL,S,_ adjusted **Eq. 7** random distribution). Normalizing these to the IEL within TEM images as for the previous two scenarios, **5 HIEL are expected per 1000µm length of IEL** (**Eq. 10**).

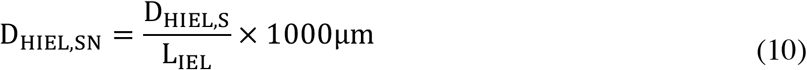

Since we have demonstrated the HIEL are randomly distributed, and there are areas sparse in HIEL, **<u>the predicted range of HIEL density in the TEM images is 5-17.5 HIEL per 1000µm IEL analyzed</u>**.

### Vascular cell co-culture

Vascular cell co-culture (VCCC) was created with human coronary artery smooth muscle cells (HCoASMCs) and human umbilical vein endothelial cells (HUVECs), paraffin embedded, and stained as originally described by us. (*6, 64*)

### Transfections for immunohistochemistry

HeLa or HEK293T were plated in 6-well dishes and grown until 70–80% confluent. Each well contained a 10mm circular glass coverslip at the bottom. Transfections were performed either with Lipofectamine 3000 or with the Lonza Nucleofector kit for HeLa cells. For validation of PS antibody, HeLa cells were plated in a 12-well dish and transfected for 17 hours with 1.8µg plasmid, 3.65µl P3000, and 2.7µl Lipofectamine 3000 per well. For validation of Kir2.1 antibody, HEK293T cells were plated in a 6-well dish and transfected for 24 hours with 2.25µg plasmid, 4.48µl P3000, and 3.47µl Lipofectamine 3000 per well. For validation of Cx40 antibody, HeLa cells were transfected using nucleofection (Lonza, VCA-1001) where 1×10^6^ cells were transfected with 0.50µg plasmid and split across two wells of a 12-well dish.

For all validation experiments, cells were fixed with 4% PFA for 10 minutes at 4°C, washed with PBS 3x 5 minutes, then blocked for 1 hour with 1% fish skin gelatin, 0.20% Triton-X 100, 0.50% BSA, and 5% animal serum in PBS. Primary antibody was diluted in blocking solution at a 1:100 concentration overnight at 4°C. The next day, following 3x 5-minute PBS washes, secondary antibody incubations were performed at a concentration of 1:500 for 1 hour at RT with gentle shaking. After secondary incubations, samples were washed 1x with PBS for 10 minutes, then incubated with DAPI at a concentration of 1:5000 for 10 minutes in PBS, and washed 1x for 10 minutes with DAPI prior to mounting the coverslips onto microscope slides. Images were obtained using an Olympus FV1000 confocal microscope.

### Western blot

Protein samples were loaded at 20µg per well and run on an 8% gel (Invitrogen, WG1001BOX) at 170V for 70 minutes. Protein was transferred to a nitrocellulose membrane (Genesee, 84-876) at 100V for 1 hour. After a 20-minute wash with diH_2_O, total protein was visualized with Revert 700 Total Protein Stain (Licor, 926-11021) and imaged on a Licor Odyssey. Total protein stain was reversed by 2x rinses with reversal solution (0.1M NaOH / 30% MeOH in diH_2_O). Next, the membrane was blocked for 1 hour in 5% BSA / TBS at RT with gentle rocking. Primary antibodies were applied at a 1:1000 dilution in blocking solution overnight at 4°C with gentle rocking. The following day, after 2x 10-minute 0.02% Tween 20 / TBS washes with medium shaking, a secondary antibody (Licor, 926-32210) was applied at a 1:10,000 dilution in blocking solution for 1 hour at RT with gentle shaking. Membranes were imaged on a Licor Odyssey. Quantification was done relative to total protein and normalized to the average value of controls.

### Whole-cell patch-clamp electrophysiology

HEK293T cells at 60% confluence (10cm dish) were transfected for 24 hours with 14µg of pCAG-Kir2.1-T2A-tdTomato (Addgene, 60598) using 21.7µl Lipofectamine 3000 (Thermofisher, L3000008) and 28µl P3000 (Thermofisher, L3000008). Transfection was confirmed prior to experiment by td-Tomato fluorescence using a confocal microscope epi-lamp.

Currents were recorded in a conventional whole cell patch clamp configuration with a voltage ramp from -140 mV to 50 mV over 250 ms, and were measured with 10µM diC8PIP_2_ (10mM stock in Krebs-HEPES, Cayman Chemical, 64910), 10µM PS (10.15 mg/ml stock in 1:1 EtOH:diH_2_O, Avanti, 840035), or 10µM diC8PIP_2_ + 10µM PS in pipette solution. For each of these conditions, Ba^2+^-sensitive currents were measured by adding 100µM Ba^2+^ to the bath solution and recording current 5 minutes later. For basal measurements, current was recorded after equilibration. The pipette solution consisted of 10 mM HEPES, 30 mM KCl, 10 mM NaCl, 110 mM K-aspartate and 1 mM MgCl2 (adjusted to pH 7.2 with NaOH). HEPES-PSS was used as the bath solution (10 mM HEPES, 134 mM NaCl, 6 mM KCl, 1 mM MgCl_2_ hexahydrate, 2 mM CaCl_2_ dihydrate, and 7 mM dextrose, pH adjusted to 7.4 using 1 M NaOH).

Patch electrodes were pulled with a Narishige PC-100 puller (Narishige International USA, Inc, Amityville, NY) and polished using a MicroForge MF-830 polisher (Narishige International USA). The pipette resistance was (3–5 ΩM). Data were acquired using a Multiclamp 700B amplifier connected to a Digidata 1550B system and analyzed using Clampfit 11.1 software (Molecular Devices, San Jose, CA, USA).

### General Pressure Myography

For all experiments, Krebs-HEPES buffer containing (in mM) NaCl 118.4, KCl 4.7, MgSO_4_ 1.2, NaHCO_3_ 4, KH_2_PO_4_ 1.2, HEPES 10, and glucose 6. On the day of the experiment, CaCl_2_ is added to the buffer at a final concentration of 2mM. Buffer is pH measured on day of experiment and adjusted to be within 7.40-7.42 by adding NaOH or HCl. Mice are sacrificed using CO_2_ asphyxiation. The mesentery was dissected out and placed in ice-cold buffer, and then pinned out on a 10cm plate filled with ∼0.5cm of Sylgard (Electron Microscopy Sciences, 24236-10; Fine Science Tools, 26002-10). Third order mesenteric arteries, defined as the third branch point relative to the feed artery and with a maximum diameter between 100-200µm, were cleared of surrounding adipose tissue using super fine forceps (Fine Science Tools, 11252-00) and microscissors (Fine Science Tools, 15000-03). The artery was then cut out of the mesentery, placed into the arteriograph chamber (DMT) containing Krebs-HEPES, and cannulated on glass pipette tips using super fine forceps (Fine Science Tools, 11254-20) and suture (DMT, Nylon, P/N 100115). Glass pipette tips were created from glass rods (World Precision Instruments, Inc., 1B120F-4) using a Narshige pipette puller (Model PC-100) with settings A=65 and B=55. Buffer was gently pushed through the artery to clear remaining blood before cannulating the second side. Following cannulation, the artery was equilibrated over a 25-minute period by increasing the pressure from 20 to 80 mmHg in 20 mmHg increments (Big Ben Sphygmomanometer) and slowly heating the artery to 37°C. For experiments on *Eln*^fl/fl^/Cre^-^ or *Eln*^fl/fl^/Cre^+^ mice, arteries were equilibrated to 60mmHg. The buffer was circulated between the arteriography chamber and a beaker containing an excess reservoir of buffer by using a peristaltic pump (Atalyst Masterflex, FH30) to prevent overheating and facilitate the delivery of pharmacological agents. DMT cell culture pressure arteriography setups were used and the inner diameter of the vessel was recorded with the 2015 release of the DMT software.

For myogenic tone experiments, third order arteries were cannulated, pressurized to 60mmHg, and equilibrated to 37°C. After the development of myogenic tone, EC function was evaluated using 1-2µM NS309. Maximum diameter was recorded 5 minutes later and NS309 was washed out of the system until myogenic tone returned. The active curve was completed by increasing the pressure in 20mmHg increments from 20-120mmHg. Diameter was recorded after 5 minutes of plateaued diameter at each pressure (D_ACT_). After the active curve, pressure was brought back down to 60mmHg for 5 minutes. SMC and EC function were evaluated with 10µM PE dose, 10µM Ach, and 30mM KCl. Next, Ca^2+^-free Krebs-HEPES was circulated in the system for 15 minutes. The passive curve was then completed by increasing the pressure in 20mmHg increments from 20-120mmHg. Diameter was recorded after 5 minutes equilibration at each pressure (D_PASS_). Myogenic tone was calculated using **Eq. 11**.

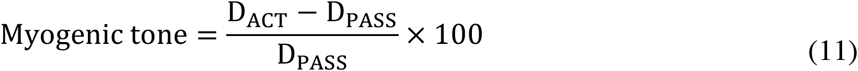

The percent change in diameter relative to 20mmHg was calculated for active and passive curves using **Eq. 12**, where D_X_ is diameter at any pressure and D_20mmHg_ is the diameter at 20mmHg.

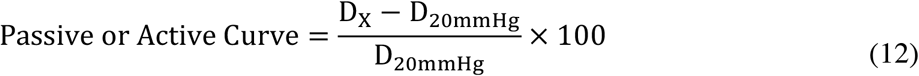

### NS309 dose response curves

For NS309 curves, the total system volume was 50ml to facilitate delivery of small doses of NS309. The artery was pre-constricted using 1µM phenylephrine (PE) in water (Sigma, P6126) and the inner diameter was recorded after 10 minutes (D_PE_). A 10mM NS309 (Sigma, N8161) in DMSO stock solution was prepared and 10µl aliquots were stored at -20°C in amber microcentrifuge tubes. The stock solution was thawed prior to the experiment and the following NS309 concentrations (in µM) were tested in the 50ml system volume: 0.1, 0.3, 0.5, 0.6, 1, and 2, where the 0.1µM dose corresponds to 0.5µl of the NS309 stock (D_NS309_). The inner diameter was recorded as the average over a 7-minute time period for each dose of NS309. SMC function was assessed by adding 30mM KCl (1M stock in water), and the inner diameter was recorded as the plateau after 5-10 minutes. The bath was then replaced with Ca^2+-^free Krebs-HEPES, containing 1mM EGTA and 100µM SNP, and inner diameter was recorded after 10 minutes (D_MAX_). The data was exported to Microsoft Excel to calculate percent dilation using **Eqn 13**.

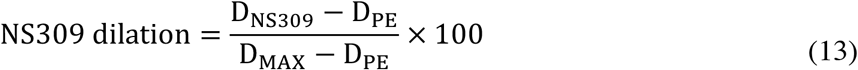

The percent vasodilation was plotted in GraphPad Prism as a dose response curve. Students t-test was performed at each dose. For experiments where Kir2.1 was inhibited, ML-133 hydrocholoride (Sigma, 422689, IC_50_ = 1.8µM) (*47*) was prepared as 7.2mM stocks in DMSO, and was added at a final concentration of 3.6µM. For Ba^2+^ experiments (Sigma, 217565), the stock solution was prepared as 100mM water and added at a final concentration of 30-100µM. For PS experiments, stock solution was prepared as a 10.15 mg/ml in 1:1 EtOH:diH_2_O (Avanti, 840035) and added at a final concentration of 10µM. The inhibitor circulated after the artery was equilibrated and before pre-constriction with PE.

### TopFluor PS experiments

Third order mesenteric arteries were dissected from male C57Bl6/J mice and secured to Sylgard squares using tungsten wire. Arteries were place in a 1.5ml Eppendorf tube containing Krebs-HEPES supplemented with 2mM Ca^2+^. Next, arteries were incubated with DAPI at 16.67 µg/ml and Alexa Fluor 647-linked hydrazide at 3.3µM in Krebs-HEPES supplemented with 2mM Ca^2+^ for 30 minutes in a 37°C water bath. The artery was then cannulated on glass canula on a pressure myograph setup. After equilibration to 80mmHg, 10µM TopFluor-PS (Avanti, 810283P) was added to the bath solution and circulated for 30 minutes. The artery was then immediately prepared *en face* and secured to a microscope slide with a droplet of Prolong Gold with DAPI and a glass coverslip (no fixation step). Arteries were imaged on an Olympus Fluoview 1000 microscope.

### Pressure myography in lipid experiments

For evaluation of PIP_2_ dilation, third order mesenteric arteries were cannulated, pressurized to 80mmHg, and equilibrated to myogenic tone. After myogenic tone plateaued and stabilized for a period of 10 minutes (D_EQ_), 10µM diC8PIP_2_ (10mM stock in Krebs-HEPES, Cayman Chemical, 64910) was added to the bath and circulated for 15 minutes in a total system volume of 10ml. The maximum change in diameter was recorded within the respective incubation period (D_PIP2_). For experiments with Kir2.1 inhibitors, drug or lipid was added after myogenic tone developed and the diameter plateaued for 10-minutes. ML-133 and PS were circulated for 30 minutes, while Ba^2+^ was circulated for 15 minutes prior to adding exogenous diC8PIP_2_. Maximum change in diameter was recorded (D_PIP2_). Next, lipids and inhibitors were washed out of the system for 5 minutes. Arterial function was then evaluated by dilation to 1µM NS309 and constriction to 30mM KCl. As with other pressure myography experiments, Ca^2+^-free Krebs-HEPES was circulated through the system at the end of the experiment and diameter was recorded 10 minutes later (D_MAX_). Two zones are monitored for arterial dimeter and the average of the two measurement are reported per artery. Zones were excluded if stable myogenic tone could not be achieved for the duration of the experiment. Lastly, if observable myogenic tone constriction occurred following a transient constriction or dilation to lipids, it was not recorded as the maximum change in diameter. For all experiments, time to maximum dilation is reported as the nearest half minute. The dilation to diC8PIP_2_ was measured by the percent change before application of the lipid (**Eq. 14**).

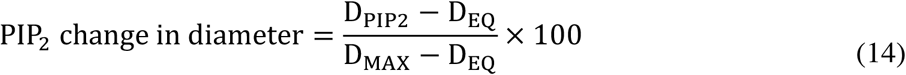

The data for maximum change in diameter for PS was taken from experiments where PS was circulating on the artery for 30 minutes prior to diC8PIP_2_ application (**Eq. 15**).

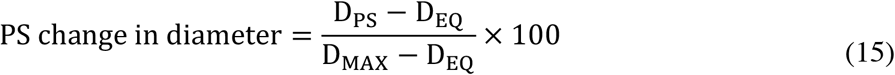

### Genomic DNA gel

Genomic DNA was isolated from lung tissue from *Eln*^fl/fl^/Cre^-^ and *Eln*^fl/fl^/Cre^+^ mice between 10-12 weeks old. Lung samples were incubated at 56°C overnight in lysis buffer (20mM Tris Base; 150mM NaCl; 1mM EDTA; 1mM EGTA; 20mM NaF; 0.5% Triton x-100) supplemented with 100ug/ml of Proteinase K. Isopropanol was used to precipitate DNA and ethanol was used to wash DNA. The dried DNA pellet was resuspended in autoclaved water and incubated for 1 hour at 56°C to allow the DNA to go into solution. The concentration was measured via nanodrop, then 15ng/ul DNA stocks were stored at -20°C until PCR was performed. The following primers were used to differentiate between wildtype and knockout mice: **WT F**: CCATGTGGGTGCTGTAAGCT, **Excision R**: GTGTGTGTAGCTGAGGAATGGG, and **LoxP Site R**: CCTACCTTTCTGGGGCCACT (ordered from Integrated DNA Technologies). Each PCR reaction contained 10µl of MyTaqRed Mix (Bioline, C755G95), 350nM of each primer, and 100ng of gDNA. Total reaction volume was 20µl. Initial denaturation temperature was 95°C for 5 minutes, followed by 28 cycles of with 30 seconds denaturation, annealing, and elongation steps at 95°C, 61°C, and 72°C, respectively. The final annealing step was 72°C for 5 minutes. PCR products were run on a 1% agarose gel containing 0.01% ethidium bromide for 50 minutes at 120V. Gels were imaged using a UV light source. The expected bands are 243 bp (WT), 283bp (KO LoxP site) and 410 bp (excision product).(49)

### qPCR

Mesenteric vasculature was trimmed of connective tissue, snap frozen in liquid nitrogen, and stored in -80°C. RNA isolation from tissue was achieved using an Aurum Total RNA Fatty and Fibrous Tissue Kit (BioRad, 7326870). RNA yields were between 40-200 ng/µl. cDNA synthesis was next performed using SuperScript IV Reverse Transcriptase (Thermofisher, 18090050). Total well volume is 20µl with equal amounts of cDNA for each sample loaded in 8µl, and master mix as 12µl and containing: (1) Taqman gene expression master mix (Thermofisher, 4369016), (2) probe of interest (Taqman probe Mm00514670_m1 *Eln* FAM-MGB, Thermofisher, 4453320), and an (3) in-well control (Taqman probe Mm00437762_m1 *B2M* VIC PL, Thermofisher, 4448485). Each sample was loaded in triplicate. Protocol was run using a Biorad Thermal Cycler using a standard protocol for 40 cycles. ΔΔC_T_ quantification method. Student’s t-test.

### VVG stain and quantification

Arteries were isolated from mice and fixed with 4% PFA at 4°C overnight, embedded in 3% agarose/PBS, then sent to the UVA Research Histology Core to be embedded in paraffin wax cut into 5µm cross-sections. The cross-sections were deparaffinized by heating to 60°C for 1 hour in a glass slide holder, followed by dehydration with Histoclear (National Diagnostics, HS-200) and a decreasing ethanol gradient. The Elastic Stain Kit (Verhoeff Van Gieson / EVG Stain) kit was purchased from Abcam (ab150667). Slides were placed into elastic stain solution for 15 minutes and rinsed with cool, running diH_2_O to remove excess stain. Sections were differentiated by dipping the slides in 2% ferric chloride solution 15 times. The differentiation reaction was stopped by rinsing slides in cool, running diH_2_O. Excess stain was further removed by dipping the slides in the sodium thiosulfate solution for 1 minute, followed by rinsing in cool, running diH_2_O. The slides were then placed in Van Gieson solution for 2 minutes for a counterstain. Excess stain was removed by rinsing in two changes of 95% EtOH, followed by a 2-minute incubation in 100% EtOH. The slides were then quickly mounted using VectaMount media and a coverslip. Images were taken using a traditional light microscope and a 20x or 40x objective.

ImageJ was used for image analysis. The stained arterial cross-section image was opened in ImageJ. The colors of the original image were then separated by “H PAS stain” color deconvolution (Image → Color → Color Deconvolution → H PAS). The indigo image was analyzed because it specifically contained the Verhoeff elastin stain. The indigo imaged was converted to a black and white mask, and the threshold of indigo was automatically determined by the ImageJ software (Process → Binary → Convert to Mask). Student’s t test.

### En face quantification for hydrazide *Eln*^fl/fl^/Cre^-^ and *Eln*^fl/fl^/Cre^+^ mice

ImageJ was used for image analysis. The hydrazide channel was projected as a Z-stack image and brightness and contrast were adjusted for visualization of HIEL. Next, Analyze → Histogram was used to see the distribution of pixel intensity. The cutoff for black pixels was determined by looking at the color-coded legend for the histogram. The number of black pixels per image was then calculated as the percentage of total pixels. Subtracting this number from 100 gives the percentage of pixels containing hydrazide signal. These numbers were plotted to determine hydrazide coverage in each image. Student’s t test.

### En face quantification for PS distribution in *en face* view

ImageJ was used for image analysis. The PS channel was thresholded and “Analyze Particles” Function was used to determine the area containing signaling and the number of puncta in the image, where particle size identification was set to be 0-Infinity. The results table in ImageJ contained the number of puncta and the coverage area of puncta. These values were normalized using the average value of control samples.

For HeatMap Analysis, the Royal Lookup Table in ImageJ was used to differentiate between low and high intensity signal. The index separating low and high intensity pixels was determined by looking at the color-coded legend for the histogram for the last blue bin. Empty pixels were defined as index 0 and were not included in the low intensity category. Number of low and high intensity pixels were then normalized to the average value of control images.

### Statistical analysis

Data are presented as mean ± SEM unless otherwise noted. Statistical tests for each experiment are listed in figure legends and respective method sections. A p-value of p<0.05 was considered to be statistically significant.

## Acknowledgments

We thank the UVA Histology Core and Advanced Microscopy Facility, as well as Dr. Janis Burt for gifting us Cx37, Cx40, and Cx43 plasmids. We also thank Dr. Ilya Levental and Dr. Nicolas Barbera for helpful discussions. Anita Impagliazzo provided illustrations.

## Funding

NIH T32 007284 (CAR, MAL, BC)

NIH F31HL149228-01 (CAR)

HL 088554 (BEI)

NIH R56 HL152420 (JEW)

## Author contributions

Conceptualization: CAR, BEI

Methodology: CAR, RP, MAL YLC, MK, BC, PJH, SMP, SKS

Investigation: CAR, RP, BEI

Visualization: CAR, BEI

Supervision: BEI

Writing—original draft: CAR, BEI

Writing—review & editing: All authors

## Data and materials availability

All data needed to evaluate the conclusions in the paper are present in the paper and/or the Supplementary Materials.

## References

1. S. J. Ahn, I. S. Fancher, J. T. Bian, C. X. Zhang, S. Schwab, R. Gaffin, S. A. Phillips, I. Levitan, Inwardly rectifying K(+) channels are major contributors to flow-induced vasodilatation in resistance arteries. J Physiol 595, 2339–2364 (2017).

2. S. K. Sonkusare, T. Dalsgaard, A. D. Bonev, M. T. Nelson, Inward rectifier potassium (Kir2.1) channels as end-stage boosters of endothelium-dependent vasodilators. J Physiol 594, 3271–3285 (2016).

3. S. J. Ahn, I. S. Fancher, S. T. Granados, N. F. Do Couto, C. L. Hwang, S. A. Phillips, I. Levitan, Cholesterol-Induced Suppression of Endothelial Kir Channels Is a Driver of Impairment of Arteriolar Flow-Induced Vasodilation in Humans. Hypertension 79, 126–138 (2022).

4. X. Shu, C. A. Ruddiman, T. C. S. t. Keller, A. S. Keller, Y. Yang, M. E. Good, A. K. Best, L. Columbus, B. E. Isakson, Heterocellular Contact Can Dictate Arterial Function. Circ Res 124, 1473–1481 (2019).

5. J. J. Zaritsky, D. M. Eckman, G. C. Wellman, M. T. Nelson, T. L. Schwarz, Targeted disruption of Kir2.1 and Kir2.2 genes reveals the essential role of the inwardly rectifying K(+) current in K(+)-mediated vasodilation. Circ Res 87, 160–166 (2000).

6. B. E. Isakson, B. R. Duling, Heterocellular contact at the myoendothelial junction influences gap junction organization. Circ Res 97, 44–51 (2005).

7. S. L. Sandow, C. E. Hill, Incidence of myoendothelial gap junctions in the proximal and distal mesenteric arteries of the rat is suggestive of a role in endothelium-derived hyperpolarizing factor-mediated responses. Circ Res 86, 341–346 (2000).

8. K. A. Dora, S. L. Sandow, N. T. Gallagher, H. Takano, N. M. Rummery, C. E. Hill, C. J. Garland, Myoendothelial gap junctions may provide the pathway for EDHF in mouse mesenteric artery. J Vasc Res 40, 480–490 (2003).

9. S. K. Sonkusare, T. Dalsgaard, A. D. Bonev, D. C. Hill-Eubanks, M. I. Kotlikoff, J. D. Scott, L. F. Santana, M. T. Nelson, AKAP150-dependent cooperative TRPV4 channel gating is central to endothelium-dependent vasodilation and is disrupted in hypertension. Sci Signal 7, ra66 (2014).

10. M. Ottolini, K. Hong, S. K. Sonkusare, Calcium signals that determine vascular resistance. Wiley Interdiscip Rev Syst Biol Med, e1448 (2019).

11. H. Han, A. Rosenhouse-Dantsker, R. Gnanasambandam, Y. Epshtein, Z. Chen, F. Sachs, R. D. Minshall, I. Levitan, Silencing of Kir2 channels by caveolin-1: cross-talk with cholesterol. J Physiol 592, 4025–4038 (2014).

12. N. Barbera, S. T. Granados, C. G. Vanoye, T. V. Abramova, D. Kulbak, S. J. Ahn, A. L. George, Jr., B. S. Akpa, I. Levitan, Cholesterol-induced suppression of Kir2 channels is mediated by decoupling at the inter-subunit interfaces. iScience 25, 104329 (2022).

13. S. Tikku, Y. Epshtein, H. Collins, A. J. Travis, G. H. Rothblat, I. Levitan, Relationship between Kir2.1/Kir2.3 activity and their distributions between cholesterol-rich and cholesterol-poor membrane domains. Am J Physiol Cell Physiol 293, C440–450 (2007).

14. A. Rosenhouse-Dantsker, Y. Epshtein, I. Levitan, Interplay Between Lipid Modulators of Kir2 Channels: Cholesterol and PIP2. Comput Struct Biotechnol J 11, 131–137 (2014).

15. F. Dabertrand, O. F. Harraz, M. Koide, T. A. Longden, A. C. Rosehart, D. C. Hill-Eubanks, A. Joutel, M. T. Nelson, PIP2 corrects cerebral blood flow deficits in small vessel disease by rescuing capillary Kir2.1 activity. Proc Natl Acad Sci U S A 118, (2021).

16. S. B. Hansen, X. Tao, R. MacKinnon, Structural basis of PIP2 activation of the classical inward rectifier K+ channel Kir2.2. Nature 477, 495–498 (2011).

17. C. M. Lopes, H. Zhang, T. Rohacs, T. Jin, J. Yang, D. E. Logothetis, Alterations in conserved Kir channel-PIP2 interactions underlie channelopathies. Neuron 34, 933–944 (2002).

18. X. Tao, J. L. Avalos, J. Chen, R. MacKinnon, Crystal structure of the eukaryotic strong inward-rectifier K+ channel Kir2.2 at 3.1 A resolution. Science 326, 1668–1674 (2009).

19. C. L. Huang, S. Feng, D. W. Hilgemann, Direct activation of inward rectifier potassium channels by PIP2 and its stabilization by Gbetagamma. Nature 391, 803–806 (1998).

20. L. A. Biwer, E. P. Taddeo, B. M. Kenwood, K. L. Hoehn, A. C. Straub, B. E. Isakson, Two functionally distinct pools of eNOS in endothelium are facilitated by myoendothelial junction lipid composition. Biochim Biophys Acta 1861, 671–679 (2016).

21. D. R. Voelker, Biochemistry of Lipids, Lipoproteins, and Membranes (Elsevier, Amsterdam, 1996).

22. A. L. Duncan, R. A. Corey, M. S. P. Sansom, Defining how multiple lipid species interact with inward rectifier potassium (Kir2) channels. Proc Natl Acad Sci U S A 117, 7803–7813 (2020).

23. T. Yeung, G. E. Gilbert, J. Shi, J. Silvius, A. Kapus, S. Grinstein, Membrane phosphatidylserine regulates surface charge and protein localization. Science 319, 210–213 (2008).

24. W. W. L. Cheng, N. D’Avanzo, D. A. Doyle, C. G. Nichols, Dual-mode phospholipid regulation of human inward rectifying potassium channels. Biophys J 100, 620–628 (2011).

25. Z. Fan, J. C. Makielski, Anionic phospholipids activate ATP-sensitive potassium channels. J Biol Chem 272, 5388–5395 (1997).

26. S. J. Lee, S. Wang, W. Borschel, S. Heyman, J. Gyore, C. G. Nichols, Secondary anionic phospholipid binding site and gating mechanism in Kir2.1 inward rectifier channels. Nat Commun 4, 2786 (2013).

27. L. A. Biwer, B. E. Isakson, Endoplasmic reticulum-mediated signalling in cellular microdomains. Acta Physiol (Oxf) 219, 162–175 (2017).

28. L. A. Biwer, M. E. Good, K. Hong, R. K. Patel, N. Agrawal, R. Looft-Wilson, S. K. Sonkusare, B. E. Isakson, Non-Endoplasmic Reticulum-Based Calr (Calreticulin) Can Coordinate Heterocellular Calcium Signaling and Vascular Function. Arterioscler Thromb Vasc Biol 38, 120–130 (2018).

29. B. E. Isakson, A. K. Best, B. R. Duling, Incidence of protein on actin bridges between endothelium and smooth muscle in arterioles demonstrates heterogeneous connexin expression and phosphorylation. Am J Physiol Heart Circ Physiol 294, H2898–2904 (2008).

30. S. L. Sandow, C. B. Neylon, M. X. Chen, C. J. Garland, Spatial separation of endothelial small- and intermediate-conductance calcium-activated potassium channels (K(Ca)) and connexins: possible relationship to vasodilator function? J Anat 209, 689–698 (2006).

31. K. Jander, J. Greulich, S. Gonnissen, N. Ale-Agha, C. Goy, P. Jakobs, S. Farrokh, C. Marziano, S. K. Sonkusare, J. Haendeler, J. Altschmied, Extra-Nuclear Functions of the Transcription Factor Grainyhead-Like 3 in the Endothelium-Interaction with Endothelial Nitric Oxide Synthase. Antioxidants (Basel) 10, (2021).

32. M. Ottolini, Z. Daneva, Y. L. Chen, E. L. Cope, R. B. Kasetti, G. S. Zode, S. K. Sonkusare, Mechanisms underlying selective coupling of endothelial Ca(2+) signals with eNOS vs. IK/SK channels in systemic and pulmonary arteries. J Physiol 598, 3577–3596 (2020).

33. P. E. McCallinhart, L. A. Biwer, O. E. Clark, B. E. Isakson, B. Lilly, A. J. Trask, Myoendothelial Junctions of Mature Coronary Vessels Express Notch Signaling Proteins. Front Physiol 11, 29 (2020).

34. M. Ottolini, K. Hong, E. L. Cope, Z. Daneva, L. J. DeLalio, J. D. Sokolowski, C. Marziano, N. Y. Nguyen, J. Altschmied, J. Haendeler, S. R. Johnstone, M. Y. Kalani, M. S. Park, R. P. Patel, W. Liedtke, B. E. Isakson, S. K. Sonkusare, Local Peroxynitrite Impairs Endothelial Transient Receptor Potential Vanilloid 4 Channels and Elevates Blood Pressure in Obesity. Circulation 141, 1318–1333 (2020).

35. A. C. Straub, J. T. Butcher, M. Billaud, S. M. Mutchler, M. V. Artamonov, A. T. Nguyen, T. Johnson, A. K. Best, M. P. Miller, L. A. Palmer, L. Columbus, A. V. Somlyo, T. H. Le, B. E. Isakson, Hemoglobin alpha/eNOS coupling at myoendothelial junctions is required for nitric oxide scavenging during vasoconstriction. Arterioscler Thromb Vasc Biol 34, 2594–2600 (2014).

36. T. Rohacs, J. Chen, G. D. Prestwich, D. E. Logothetis, Distinct specificities of inwardly rectifying K(+) channels for phosphoinositides. J Biol Chem 274, 36065–36072 (1999).

37. K. R. Heberlein, A. C. Straub, A. K. Best, M. A. Greyson, R. C. Looft-Wilson, P. R. Sharma, A. Meher, N. Leitinger, B. E. Isakson, Plasminogen activator inhibitor-1 regulates myoendothelial junction formation. Circ Res 106, 1092–1102 (2010).

38. S. J. Stone, J. E. Vance, Phosphatidylserine synthase-1 and -2 are localized to mitochondria-associated membranes. J Biol Chem 275, 34534–34540 (2000).

39. K. Del Vecchio, R. V. Stahelin, Investigation of the phosphatidylserine binding properties of the lipid biosensor, Lactadherin C2 (LactC2), in different membrane environments. J Bioenerg Biomembr 50, 1–10 (2018).

40. M. H. Andersen, H. Graversen, S. N. Fedosov, T. E. Petersen, J. T. Rasmussen, Functional analyses of two cellular binding domains of bovine lactadherin. Biochemistry 39, 6200–6206 (2000).

41. J. Shi, C. W. Heegaard, J. T. Rasmussen, G. E. Gilbert, Lactadherin binds selectively to membranes containing phosphatidyl-L-serine and increased curvature. Biochim Biophys Acta 1667, 82–90 (2004).

42. S. K. Dasgupta, P. Guchhait, P. Thiagarajan, Lactadherin binding and phosphatidylserine expression on cell surface-comparison with annexin A5. Transl Res 148, 19–25 (2006).

43. S. Mather, K. A. Dora, S. L. Sandow, P. Winter, C. J. Garland, Rapid endothelial cell-selective loading of connexin 40 antibody blocks endothelium-derived hyperpolarizing factor dilation in rat small mesenteric arteries. Circ.Res. 97, 399–407 (2005).

44. N. Shen, J. Zheng, H. Liu, K. Xu, Q. Chen, Y. Chen, Y. Shen, L. Jiang, Y. Chen, Barium chloride impaired Kir2.1 inward rectification in its stably transfected HEK 293 cell lines. Eur J Pharmacol 730, 164–170 (2014).

45. N. Alagem, M. Dvir, E. Reuveny, Mechanism of Ba(2+) block of a mouse inwardly rectifying K+ channel: differential contribution by two discrete residues. J Physiol 534, 381–393 (2001).

46. M. Wu, H. Wang, H. Yu, E. Makhina, J. Xu, E. S. Dawson, C. R. Hopkins, C. W. Lindsley, O. B. McManus, M. Li, “A potent and selective small molecule Kir2.1 inhibitor” in Probe Reports from the NIH Molecular Libraries Program (Bethesda (MD), 2010).

47. H. R. Wang, M. Wu, H. Yu, S. Long, A. Stevens, D. W. Engers, H. Sackin, J. S. Daniels, E. S. Dawson, C. R. Hopkins, C. W. Lindsley, M. Li, O. B. McManus, Selective inhibition of the K(ir)2 family of inward rectifier potassium channels by a small molecule probe: the discovery, SAR, and pharmacological characterization of ML133. ACS Chem Biol 6, 845–856 (2011).

48. A. D. Bonev, M. T. Nelson, ATP-sensitive potassium channels in smooth muscle cells from guinea pig urinary bladder. Am J Physiol 264, C1190–1200 (1993).

49. C. J. Lin, M. C. Staiculescu, J. Z. Hawes, A. J. Cocciolone, B. M. Hunkins, R. A. Roth, C. Y. Lin, R. P. Mecham, J. E. Wagenseil, Heterogeneous Cellular Contributions to Elastic Laminae Formation in Arterial Wall Development. Circ Res 125, 1006–1018 (2019).

50. C. Wilson, X. Zhang, M. D. Lee, M. MacDonald, H. R. Heathcote, N. M. N. Alorfi, C. Buckley, S. Dolan, J. G. McCarron, Disrupted endothelial cell heterogeneity and network organization impair vascular function in prediabetic obesity. Metabolism 111, 154340 (2020).

51. F. Nakatsu, A. Kawasaki, Functions of Oxysterol-Binding Proteins at Membrane Contact Sites and Their Control by Phosphoinositide Metabolism. Front Cell Dev Biol 9, 664788 (2021).

52. J. Chung, F. Torta, K. Masai, L. Lucast, H. Czapla, L. B. Tanner, P. Narayanaswamy, M. R. Wenk, F. Nakatsu, P. De Camilli, INTRACELLULAR TRANSPORT. PI4P/phosphatidylserine countertransport at ORP5- and ORP8-mediated ER-plasma membrane contacts. Science 349, 428–432 (2015).

53. S. Mochizuki, H. Miki, R. Zhou, Y. Noda, The involvement of oxysterol-binding protein related protein (ORP) 6 in the counter-transport of phosphatidylinositol-4-phosphate (PI4P) and phosphatidylserine (PS) in neurons. Biochem Biophys Rep 30, 101257 (2022).

54. D. Ma, N. Zerangue, Y. F. Lin, A. Collins, M. Yu, Y. N. Jan, L. Y. Jan, Role of ER export signals in controlling surface potassium channel numbers. Science 291, 316–319 (2001).

55. C. Stockklausner, J. Ludwig, J. P. Ruppersberg, N. Klocker, A sequence motif responsible for ER export and surface expression of Kir2.0 inward rectifier K(+) channels. FEBS Lett 493, 129–133 (2001).

56. B. E. Isakson, S. I. Ramos, B. R. Duling, Ca2+ and inositol 1,4,5-trisphosphate-mediated signaling across the myoendothelial junction. Circ Res 100, 246–254 (2007).

57. B. Razani, J. A. Engelman, X. B. Wang, W. Schubert, X. L. Zhang, C. B. Marks, F. Macaluso, R. G. Russell, M. Li, R. G. Pestell, D. Di Vizio, H. Hou, Jr., B. Kneitz, G. Lagaud, G. J. Christ, W. Edelmann, M. P. Lisanti, Caveolin-1 null mice are viable but show evidence of hyperproliferative and vascular abnormalities. J Biol Chem 276, 38121–38138 (2001).

58. A. Fujita, J. Cheng, K. Tauchi-Sato, T. Takenawa, T. Fujimoto, A distinct pool of phosphatidylinositol 4,5-bisphosphate in caveolae revealed by a nanoscale labeling technique. Proc Natl Acad Sci U S A 106, 9256–9261 (2009).

59. G. D. Fairn, N. L. Schieber, N. Ariotti, S. Murphy, L. Kuerschner, R. I. Webb, S. Grinstein, R. G. Parton, High-resolution mapping reveals topologically distinct cellular pools of phosphatidylserine. J Cell Biol 194, 257–275 (2011).

60. J. Saliez, C. Bouzin, G. Rath, P. Ghisdal, F. Desjardins, R. Rezzani, L. F. Rodella, J. Vriens, B. Nilius, O. Feron, J. L. Balligand, C. Dessy, Role of caveolar compartmentation in endothelium-derived hyperpolarizing factor-mediated relaxation: Ca2+ signals and gap junction function are regulated by caveolin in endothelial cells. Circulation 117, 1065–1074 (2008).

61. P. G. Frank, Endothelial caveolae and caveolin-1 as key regulators of atherosclerosis. Am J Pathol 177, 544–546 (2010).

62. A. Arcangeli, A. Becchetti, Complex functional interaction between integrin receptors and ion channels. Trends Cell Biol 16, 631–639 (2006).

63. S. Sengupta, K. E. Rothenberg, H. Li, B. D. Hoffman, N. Bursac, Altering integrin engagement regulates membrane localization of Kir2.1 channels. J Cell Sci 132, (2019).

64. L. A. Biwer, C. Lechauve, S. Vanhoose, M. J. Weiss, B. E. Isakson, A Cell Culture Model of Resistance Arteries. J Vis Exp, (2017).

