## Supplemental Figures and Tables for "Polarized localization of phosphatidylserine in endothelium regulates Kir2.1"

### Supplemental Materials for
### Polarized localization of phosphatidylserine in endothelium regulates Kir2.1

Claire A. Ruddiman, Richard Peckham, Melissa A. Luse, Yen-Lin Chen, Maniselvan Kuppusamy, Bruce Corliss, P. Jordan Hall, Chien-Jung Lin, Shayn M Peirce, Swapnil K. Sonkusare, Robert P. Mecham, Jessica E. Wagenseil, Brant E. Isakson*

**This PDF file includes:**

Figs. S1 to S8
Tables S1 to S3

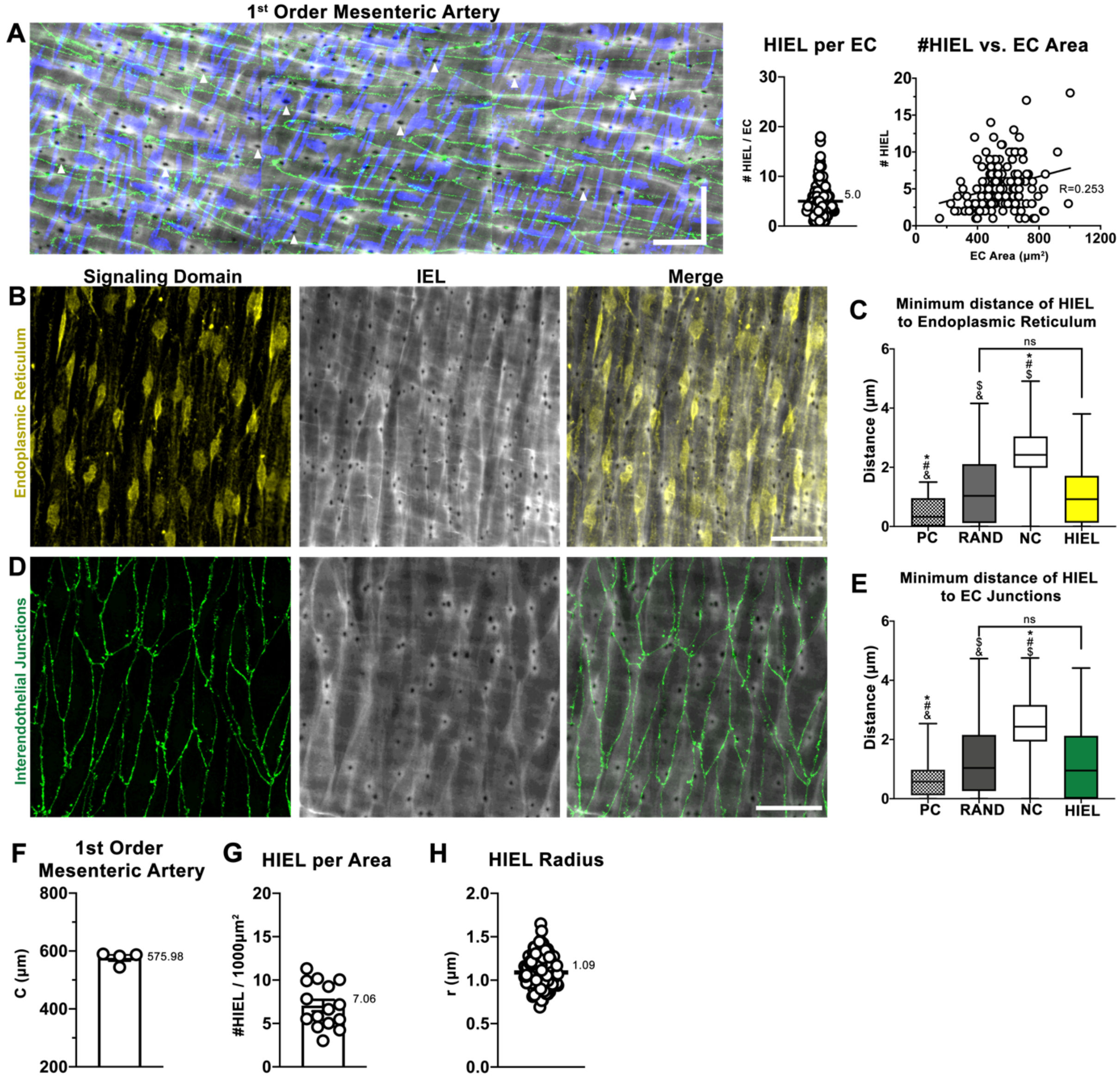


**Figure S1: HIEL are randomly distributed with respect to endothelial signaling hubs in first order mesenteric arteries. (A)** Representative stitched confocal image of a first order mesenteric artery prepared *en face* and stained for nuclei (blue) via DAPI, the IEL (grey) via Alexa-Fluor 488-linked hydrazide, and interendothelial junctions (green) via claudin-5. Scale bar is 30µm in both directions. Quantification of HIEL per EC and correlation of HIEL per EC versus EC area. N=4 mice, n=4 arteries, n=12 ROIs, and n=155 ECs. (**B**) Representative *en face* confocal image of endoplasmic reticulum (ER, yellow) detected via calnexin and IEL (grey), (**C**) box and whiskers plot of minimum distance of real-world HIEL centers to ER compared to Matlab-simulated HIEL centers. N=1 mouse, n=1 artery, n=2 ROIs, Area= $6.96 \times 10^4$ µm$^2$, and n=315 HIEL. (**D**) Representative *en face* confocal image of interendothelial junctions (green) detected via claudin-5 and IEL (grey), (**E**) box and whiskers plot of minimum distance of real-world HIEL centers to interendothelial junctions compared to Matlab-simulated HIEL centers. N=6 mice, and n=10 arteries, n=15 ROIs, Area=$1.66 \times 10^5$ µm$^2$, and n=1200 HIEL. Statistical test: Brown-Forsythe and

Welch ANOVA. # indicates a p <0.0001 significant difference to real-world HIEL distribution, * indicates a p <0.0001 significant difference to RAND, $ indicates a p <0.0001 significant difference to NC distribution, and & indicates a p <0.0001 significant difference to PC. (**F**) Circumference (C) of first order mesenteric arteries measured as the width of the artery in an *en face* preparation. N=4 mice, n=4 arteries. (**G**) HIEL per area taken from *en face* images that were used for HIEL spatial pattern analysis with respect to claudin-5. Number of HIEL was determined via the in-house Matlab program. (**H**) HIEL radius measurements obtained from in-house Matlab program.

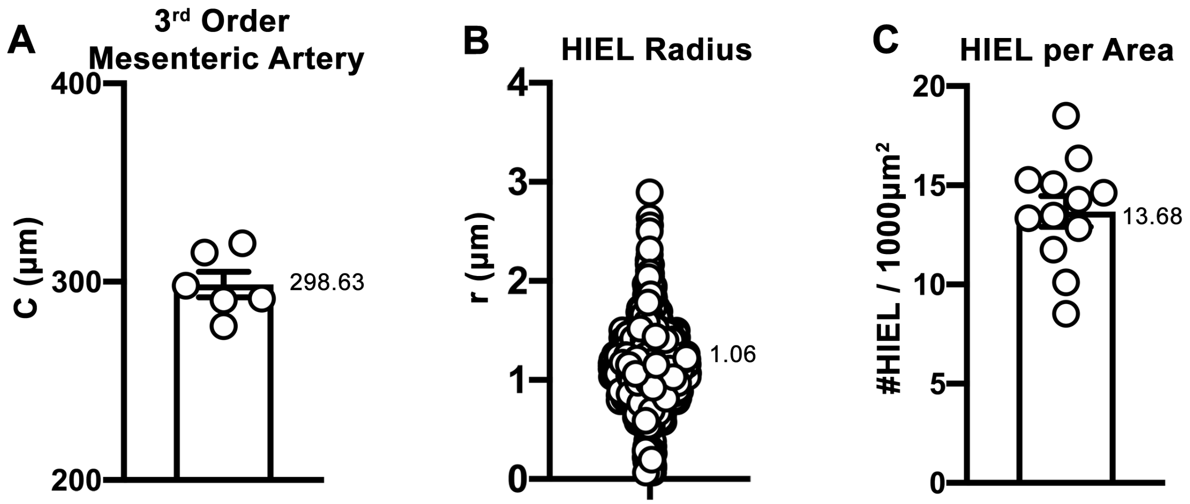


**Figure S2: Quantitative data obtained from third order mesenteric artery *en face* preparations.** (**A**) Circumference of third order mesenteric arteries measured as the width of the artery in an *en face* preparation. N=5 mouse, n=6 arteries. (**B**) HIEL radius measurements obtained from in-house Matlab program. N=6 mice, and n=10 arteries, n=22 ROIs, Area=$1.48x10^5 \mu m^2$, and n=2166 HIEL. (**C**) HIEL per area taken from *en face* images used in Matlab analysis. Number of HIEL was determined via the in-house Matlab program. N=4 mice, n=4 arteries, n=12 ROIs, and Area=$1.67x10^5 \mu m^2$.

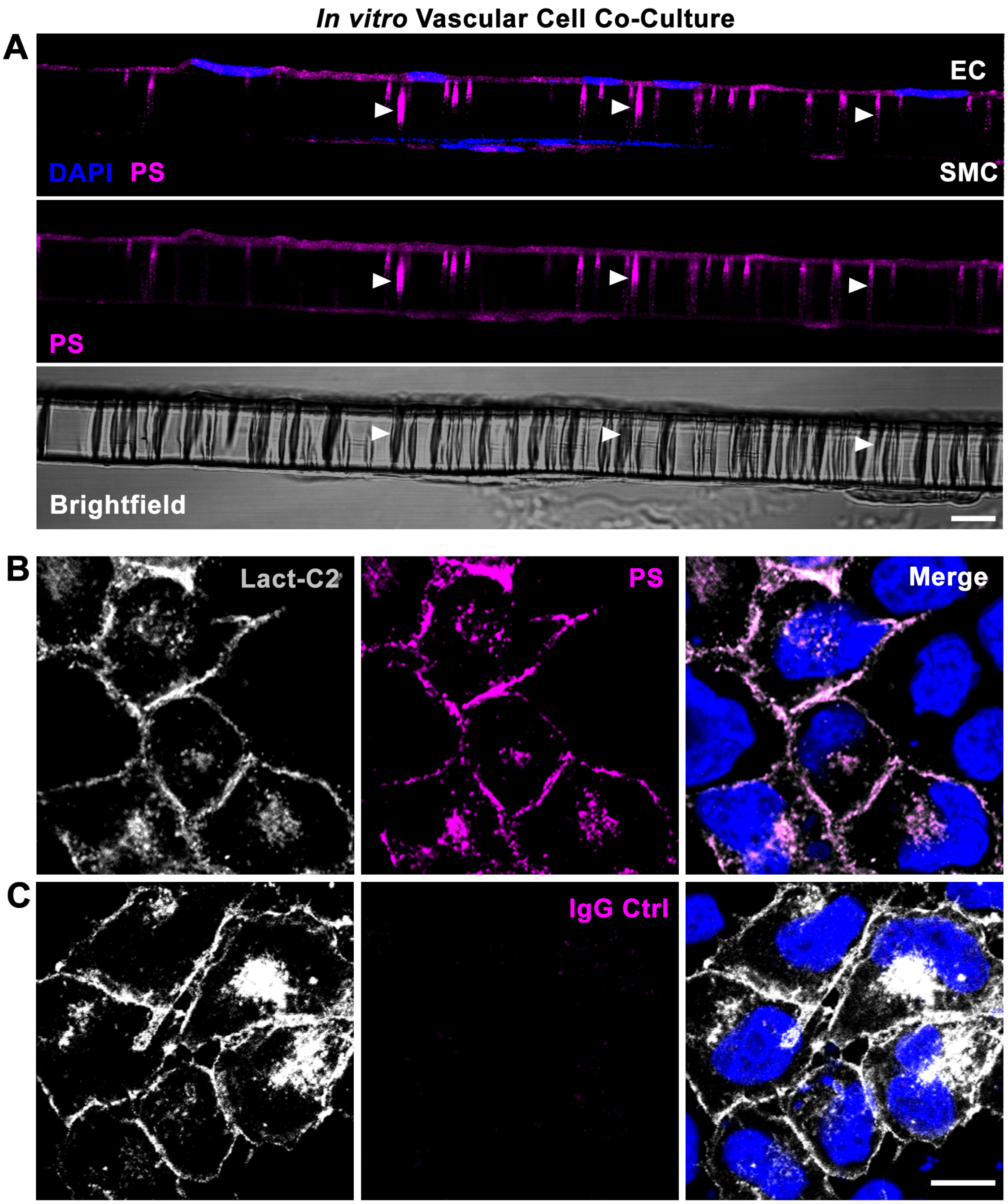


**Figure S3: Validation of PS antibody. (A)** Transverse cross-sections of an *in vitro* vascular cell co-culture model where EC and SMC are plated on either side of a Transwell. Nuclei (blue) are detected via DAPI and PS (magenta). Brightfield image of Transwell is shown in the bottom panel. Arrowheads indicate PS localization to *in vitro* MEJs. (**B**) HeLa cells transfected with Lact-C2-GFP plasmid (white) and co-stained with PS antibody or (**C**) IgG control. Nuclei (blue) are detected via DAPI. Scale bars are 10μm.

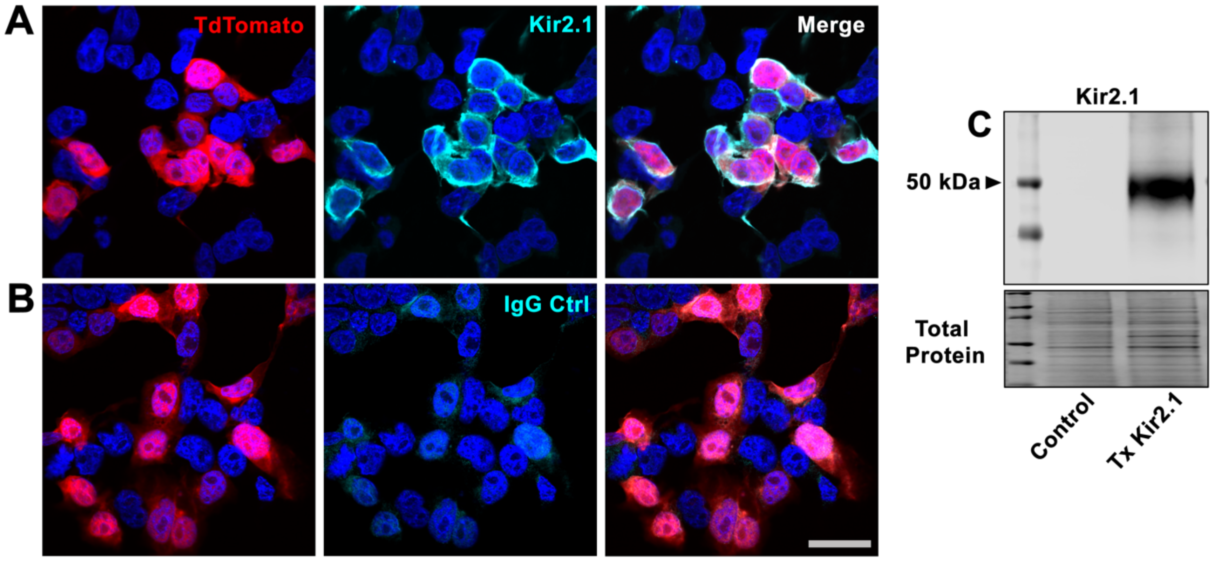


**Figure S4: Validation of Kir2.1 antibody.** (**A**) HEK293T cells transfected with Kir2.1-T2A-tdTomato plasmid (red) and co-stained with PS antibody or (**B**) IgG control. Nuclei (blue) are detected via DAPI. Scale bar is 30µm. (**C**) Western blot detection of Kir2.1 protein in HEK293T cells that were untreated or transfected with Kir2.1-T2A-tdTomato plasmid. Total protein was used as a loading control.

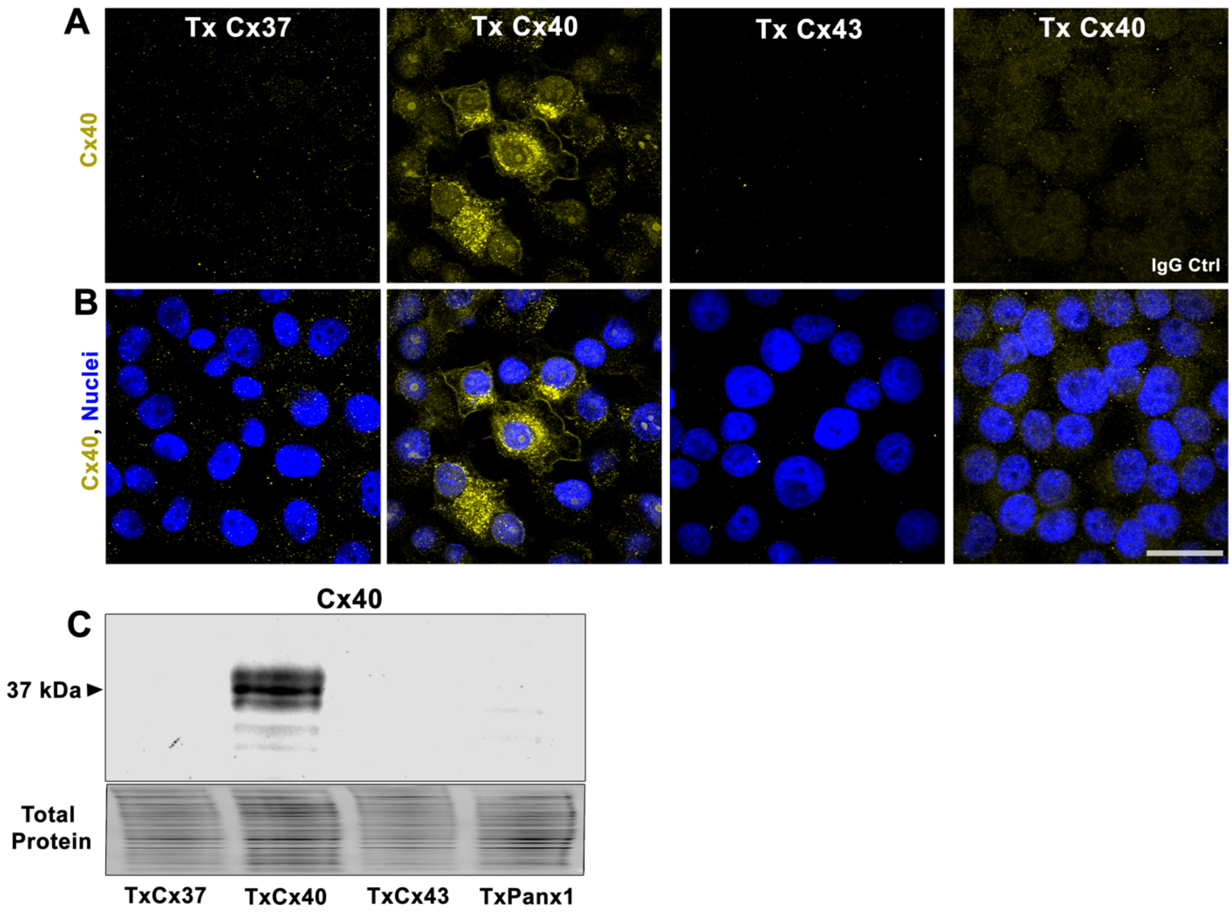


**Figure S5: Validation of Cx40 antibody.** (**A**) HeLa cells transfected with a connexin plasmid, either connexin 37 (Tx Cx37), connexin 40 (Tx Cx40), or connexin 43 (Tx Cx43), then stained using a Cx40 antibody or IgG control to evaluate Cx40 antibody specificity. Nuclei (blue) are detected via DAPI. Scale bar is 30µm. (**C**) Western blot detection of Cx40 protein in HeLa cells that were transfected with plasmids for Cx37, Cx40, Cx43, and Panx1. Total protein was used as a loading control.

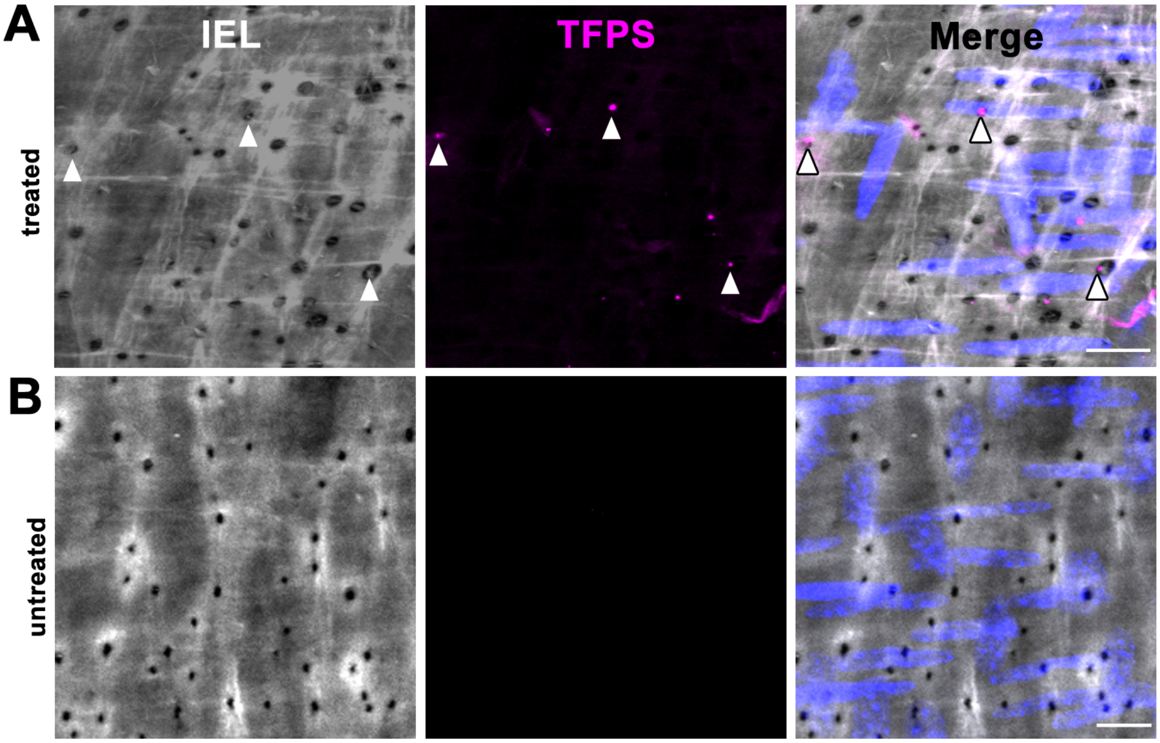


**Figure S6: Exogenous application of TopFluor-PS localizes to the MEJ in intact third order mesenteric arteries.** Live third order mesenteric arteries prepared *en face* after treatment with (**A**) 10µM TopFluor-PS in the bath solution of a pressure myography setup or (**B**) without treatment. Nuclei (blue) are detected via DAPI and IEL (grey) is detected via Alexa Fluor linked hydrazide. Arrowheads indicate TopFluor-PS localization to MEJ. Scale bars are 10µm.

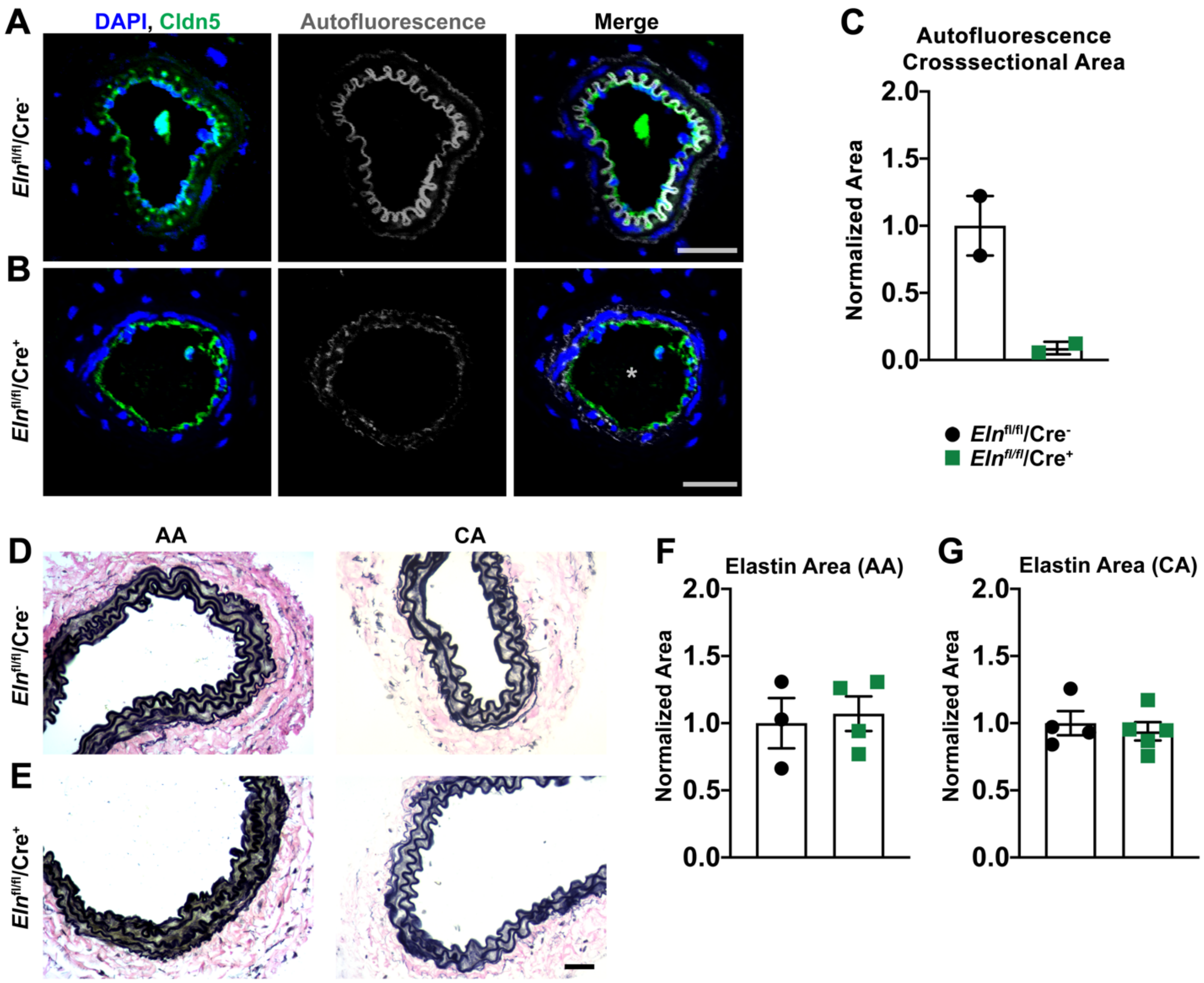


**Figure S7: EC-specific knockout of elastin disrupts IEL in resistance arteries and not large conduit arteries.** Representative images of cross-sections from third order mesenteric arteries taken from (**A**) *Eln*$^{fl/fl}$/Cre$^-$ and (**B**) *Eln*$^{fl/fl}$/Cre$^+$ mice where nuclei (blue) are detected via DAPI, IEL (grey) is detected via autofluorescence, and interendothelial junctions (green) are detected via claudin-5. Scale bar is 30µm. N=2 mice per group. (**C**) Quantification of cross-sectional IEL area detected via autofluorescence. Representative images of cross-sections from the abdominal aorta (AA) and carotid artery (CA) of (**D**) *Eln*$^{fl/fl}$/Cre$^-$ and (**E**) *Eln*$^{fl/fl}$/Cre$^+$ mice. Scale bar is 30µm. Quantification of Verhoeff stain in (**F**) AA and (**G**) CA. N= 3-5 mice per group. Student's t-test.

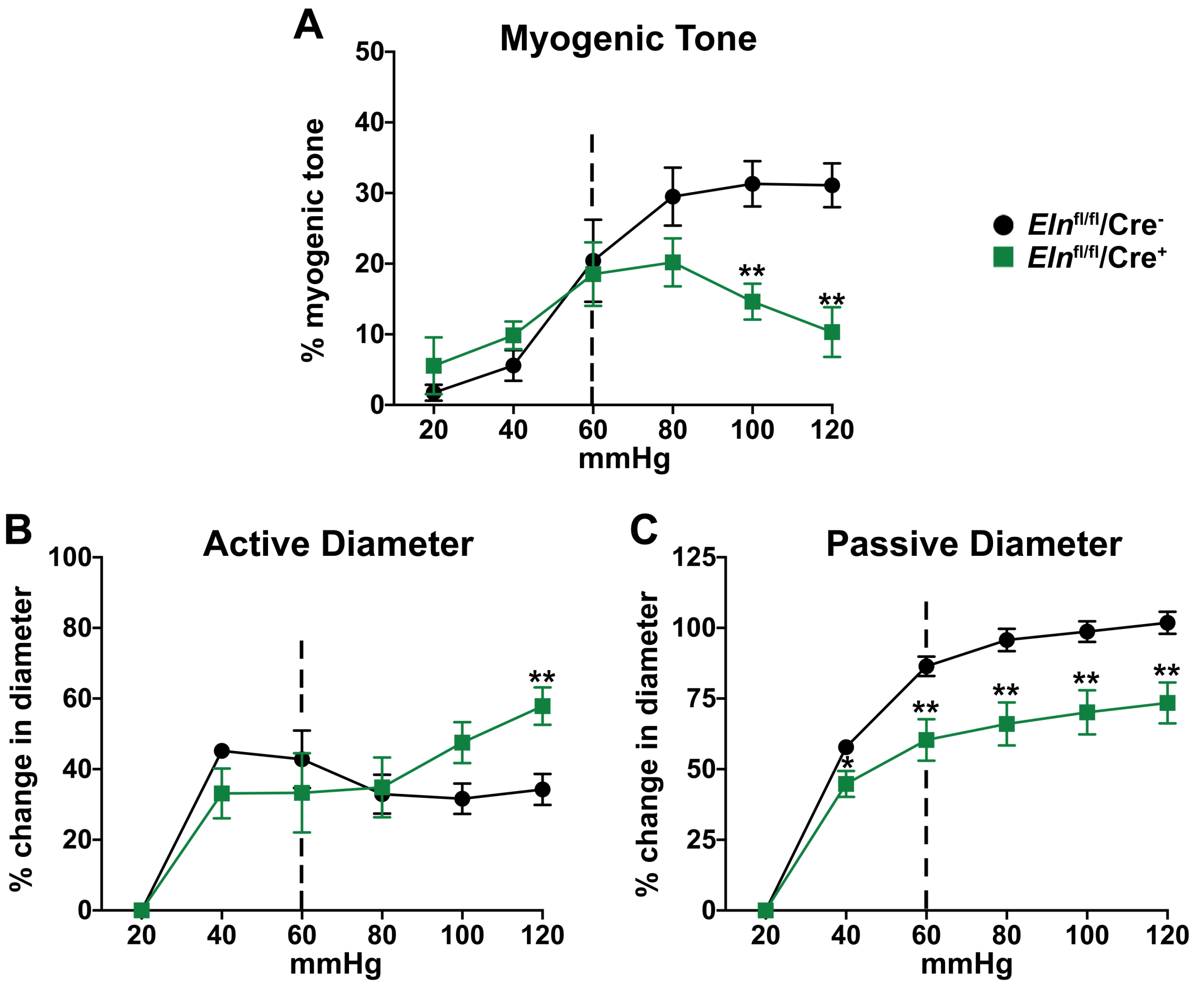


**Figure S8: Determination of optimal pressure for myography experiments on *Eln*$^{fl/fl}$/Cre$^{+}$ mice.** Pressure myography experiments on third order mesenteric arteries from *Eln*$^{fl/fl}$/Cre$^{-}$ and *Eln*$^{fl/fl}$/Cre$^{+}$ mice, where (**A**) is myogenic tone, (**B**) is the active diameter, and (**C**) is the passive diameter. Dotted line indicates the optimal pressure for *Eln*$^{fl/fl}$/Cre$^{+}$ arteries. N=3 mice per group, n=4-6 arteries per group. Students t-test was performed at each pressure. * indicates $p<0.050$, and ** indicates $p<0.010$.

| Variable | Value | Unit | Source | Description |
|---|---|---|---|---|
| **C** | 298.6 | µm | N=4, n=6 arteries **Fig. S2A** | Circumference of 3rd order mesenteric arteries |
| **$C_{IEL}$** | 298.6 | µm | Assume that it is equal to C | Circumference of IEL |
| **d** | 2.1 | µm | N=5 mice, n=2166 HIEL **Fig. S2B** | Average diameter of HIEL |
| **$\bar{A}_{xy}$** | 13,942.9 | $\mu m^2$ | Experimental parameter | Area of en face images |
| **$\tilde{N}_{HIEL}$** | 190.7 | HIEL / image | N=4, n=4 arteries, n=12 images **Fig. S2C** | Average number of HIEL |
| **$\rho_{HIEL}$** | 13.6 | HIEL per $1000\mu m^2$ | **Eq. 1** | Density of HIEL per $1000\mu m^2$ |

**Table S1: Summary of quantitative data obtained from *en face* images.** Measurements taken from third order mesenteric arteries, either manually, automatically via in-house Matlab program, or calculated from direct measurements.

| Variable | Value | Units | Source | Description |
|---|---|---|---|---|
| $\mathbf{Y_{TEM}}$ | 0.070 | µm | Experimental parameter | Thickness of transverse TEM sections |
| $\mathbf{A_{TEM,1}}$ | 20.902 | $\mu m^2$ | **Eq. 2** | Artery area spanning $\mathbf{Y_{TEM}}$ |
| $\mathbf{A_{TEM,\,d}}$ | 627.06 | $\mu m^2$ | **Eq. 4** | Artery area spanning the length **d** |
| $\mathbf{L_{IEL}}$ | 1791.6 | µm | **Eq. 5** | Length of IEL across 6 TEM sections |
| $\mathbf{HIEL_{TEM}}$ | 9 | HIEL per $A_{TEM}$ | **Eq. 6** | Number of HIEL expected across $A_{TEM,d}$ |

**Table S2: Calculated parameters of arterial cross-sections imaged via TEM.** Experimental and calculated parameters for converting *en face* quantitative data to be interpretable in a TEM geometry.

| Case Definition | Variable | Value | Units | Equation | Description |
|---|---|---|---|---|---|
| **Case1: Maximum case, assumes HIEL are evenly distributed** | $D_{HIEL, E}$ | 54 | TEM detections | **Eq. 7** | HIEL detections over 6 sections for Case 1 |
| **Case 2: Assumes HIEL are randomly distributed** | $D_{HIEL, R}$ | 32 | | **Adjusted Eq. 7 (see text)** | HIEL detections over 6 sections for Case 2 |
| **Case 3: Minimum case, represents the rare case where each HIEL only appear on 1/6 of sections** | $D_{HIEL, S}$ | 9 | | **Adjusted Eq. 7 (see text)** | HIEL detections over 6 sections for Case 3 |
| **Normalized Case 1** | $D_{HIEL, EN}$ | 30 | TEM detections per 1000μm IEL | **Eq. 8** | HIEL detections over 6 sections for Case 1 normalized to IEL length |
| **Normalized Case 2** | $D_{HIEL, RN}$ | 17.8 | | **Eq. 9** | HIEL detections over 6 sections for Case 2 normalized to IEL length |
| **Normalized Case 3** | $D_{HIEL, SN}$ | 5 | | **Eq. 10** | HIEL detections over 6 sections for Case 3 normalized to IEL length |

**Table S3: Case scenarios that consider the potential spatial distributions of HIEL.** Prediction of HIEL incidence in TEM sections with consideration of spatial distributions of evenly distributed (**Case 1**), randomly distributed (**Case 2**), or sparsely distributed (**Case 3**). Predictions are then normalized to obtain ratios with respect to IEL length, a metric that can be reproducibly measured in TEM images (**Normalized Cases 1-3**).
